# Lacustrine speciation associated with chromosomal inversion in a lineage of riverine fishes

**DOI:** 10.1101/2022.12.12.519811

**Authors:** Daniel J. MacGuigan, Trevor J. Krabbenhoft, Richard C. Harrington, Dylan K. Wainwright, Nathan J. C. Backenstose, Thomas J. Near

**Author notes:** **Author Contributions:** D.J.M., T.J.K., and T.J.N. designed research; D.J.M., T.J.K., R.C.H., D.K.W., N.J.C.B, and T.J.N. performed research; D.J.M., T.J.K., R.C.H., D.K.W., N.J.C.B, and T.J.N. contributed new reagents/analytic tools; D.J.M., T.J.K., R.C.H., and D.K.W. analyzed data; D.J.M., T.J.K., R.C.H., D.K.W., and T.J.N. wrote the paper. **Data Availability:** The *Etheostoma perlongum* genome assembly is available through NCBI (PRJNA682188) and all associated sequence data (Nanopore, 10x Chromium, Hi-C Illumina reads, and RNA-seq) are deposited in the NCBI SRA (PRJNA682188). The genome annotation will be made available through Dryad. The *Etheostoma nigrum* genome assembly is also available through NCBI (PRJNA895006) and all associated sequence data (Nanopore, Hi-C Illumina reads) are deposited in the NCBI SRA (SRR22084753). Demultiplexed ddRAD sequence data are available on NCBI (PRJNA835500). Sequence data will be released upon publication or will be provided upon editor/reviewer request. Meristic data, morphometric data, osteological measurements, and diet will be made available through Dryad upon publication and is available upon editor/reviewer request. Images used for 2D body shape morphometrics and TPS landmark data will be made available through Dryad. All 3D models generated from µCT scans will be made accessible through MorphoSource upon publication (https://www.morphosource.org/). **Code Availability:** Code for genome assembly and analyses will be made available through Dryad upon publication and is available upon editor/reviewer request.

## Abstract

Geographic isolation is the primary driver of speciation in many vertebrate lineages. This trend is exemplified by North American darters, a clade of freshwater fishes where nearly all sister species pairs are allopatric and separated by millions of years of divergence. One of the only exceptions is the Lake Waccamaw endemic *Etheostoma perlongum* and its riverine sister species *E. maculaticeps,* which have no physical barriers to gene flow. Here we show that lacustrine speciation of *E. perlongum* is characterized by morphological and ecological divergence likely facilitated by a large chromosomal inversion. While *Etheostoma perlongum* is phylogenetically nested within the geographically widespread *E. maculaticeps*, there is a sharp genetic and morphological break coinciding with the lake-river boundary in the Waccamaw River system. Despite recent divergence, an active hybrid zone, and ongoing gene flow, analyses using a *de novo* reference genome reveal a 9 Mb chromosomal inversion with elevated divergence between *E. perlongum* and *E. maculaticeps.* This region exhibits striking synteny with known inversion supergenes in two distantly related fish lineages, suggesting deep evolutionary convergence of genomic architecture. Our results illustrate that rapid, ecological speciation with gene flow is possible even in lineages where geographic isolation is the dominant mechanism of speciation.

## INTRODUCTION

Geographic isolation historically dominated discussion of speciation mechanisms^1^. For instance, Ernst Mayr largely dismissed the idea that environmental breaks could lead to speciation, stating “no case is known to us that could be cited as irrefutable proof for…[ecological] speciation”^2^. This mindset percolated into many subdisciplines of biology, especially freshwater ichthyology where some concluded that there are no known examples of parapatric or sympatric speciation that show an “ecological mode”, in part because “[ichthyologists] tend to think ‘allopatrically’”^3^. However, a large body of theoretical and empirical literature demonstrates that ecological speciation without geographic isolation is possible, particularly in lakes where heterogeneous and highly structured environments can drive adaptive diversification^4^. Lakes serve as biogeographic “islands” that geographically and ecologically isolate emerging species from nearby freshwaters, such as an endemic minnow in Lake Biwa, Japan^5^, a species complex of ricefishes in Sulawesi^6^, sailfin silversides in Lake Matano, Indonesia^7^, and Arctic Charr in many postglacial lakes^8^. Despite myriad ecological speciation studies in lakes, taxonomy often lags behind. For instance, many lake-stream pairs of threespine sticklebacks are in the early stages of parapatric speciation and the diverse stickleback morphotypes are often considered a “complex of species”^9^, yet no taxonomic revision has ever been proposed. Discrepancy between speciation studies and taxonomy can have conservation consequences and lead to underestimates of lacustrine biodiversity^10^.

One of the rare cases where taxonomy has outpaced speciation research is Lake Waccamaw, one of the largest Carolina Bay lakes that exemplifies the insular nature of lacustrine ecosystems. Lake Waccamaw is a shallow, sandy lake (max depth 3.3 m), with neutral pH, high alkalinity, clear waters, and open habitat compared to the surrounding acidic, highly structured blackwater habitats of the Waccamaw River basin^11–14^. Although Lake Waccamaw is only 15,000-32,000 years old^14^, it contains three phylogenetically disparate endemic fish species: the Waccamaw Silverside (*Menidia extensa*), the Waccamaw Killifish (*Fundulus waccamensis*), and the Waccamaw Darter (*Etheostoma perlongum*)^15^, as well as two endemic mussels and an endemic snail^16, 17^. Morphological differences between the Lake Waccamaw endemics and their closest riverine relatives suggest lacustrine adaptation. Each lake species has more vertebrae, more lateral line scales, and slender, elongated body shapes^15, 18^. Perhaps most unusual among the Lake Waccamaw endemics is the Waccamaw Darter, *Etheostoma perlongum*. Of the approximately 250 species of darters, 93% of sister species pairs are allopatric^19^, yet, *E. perlongum* is not geographically isolated from its presumed sister lineage, the riverine *Etheostoma maculaticeps*^20^. Most darter species exclusively occupy riverine habitats and *E. perlongum* is the only entirely lacustrine species. Although some darter species exhibit intraspecific lake-river morphological differences^21^, *E. perlongum* is the only potential case of lacustrine speciation in the clade. Lack of evidence for strong genetic differentiation and morphological variation within the Waccamaw River basin has led to questions about the distinctiveness of *E. perlongum* as a species^22, 23^.

In this study, we investigate the ecological, morphological, and genomic context of speciation between *Etheostoma perlongum* in Lake Waccamaw and *E. maculaticeps* in the adjacent Waccamaw River. We assemble and annotate a *de novo* chromosome-level genome for *E. perlongum* and analyze thousands of double digest restriction site associated DNA (ddRAD) markers from individuals spanning a lake-river transect. Using this rich genomic dataset, we determine if there are regions of the genome resistant to gene flow that potentially reveal the genomic architecture of speciation between *E. perlongum* and *E. maculaticeps*. We complement genomic analyses with morphometric analyses of body shape, analyses of ecomorphological osteology with microcomputed tomography (µCT) imaging, and diet analysis of stomach contents. Our analyses reveal genomic, morphological, and ecological differentiation between darters from Lake Waccamaw and Waccamaw River. We detect substantial, ongoing gene flow between *E. perlongum* and *E. maculaticeps* facilitated by an active hybrid zone in the Waccamaw River just downstream of Lake Waccamaw. Despite gene flow, we find elevated genomic divergence between *E. perlongum* and *E. maculaticeps* localized in a 9 Mb region of chromosome 9 with high levels of linkage disequilibrium, likely the result of a chromosomal inversion. Gene annotations suggest that this inversion is a supergene involved in retinal development, olfactory signaling, and circadian regulation. We hypothesize that the chromosome 9 inversion is subject to differential selective pressures in lake and river environments driving ecological and morphological divergence. Our results reveal that in a clade where diversification is primarily the product of geographic isolation, modifications to genomic architecture can facilitate ecological divergence and speciation in the face of gene flow.

## MATERIALS AND METHODS

### Sampling and ddRAD Sequencing

We collected tissue samples for 93 individuals along a transect encompassing five sites in Lake Waccamaw and seven sites in the Waccamaw River basin (Fig. 1A, Table S1). In addition, we sampled 43 individuals from 28 localities across the range of *Etheostoma maculaticeps*, the closest relative of *E. perlongum*, and two individuals from each of two outgroup species, *E. olmstedi* and *E. nigrum* (Table S1). Following Yale IACUC protocol number 2018-10681, fish were euthanized by immersion in 250 mg/L buffered MS-222 solution before collecting tissue samples, typically from the right pectoral fin. Tissues were preserved in 95% ethanol at −20 °C. We performed a modified version of a double digest restriction site associated DNA (ddRAD)^24, 25^. Details of DNA extraction, library preparation, and assembly may be found in the Supporting Information and Table S2.

**Figure 1.**
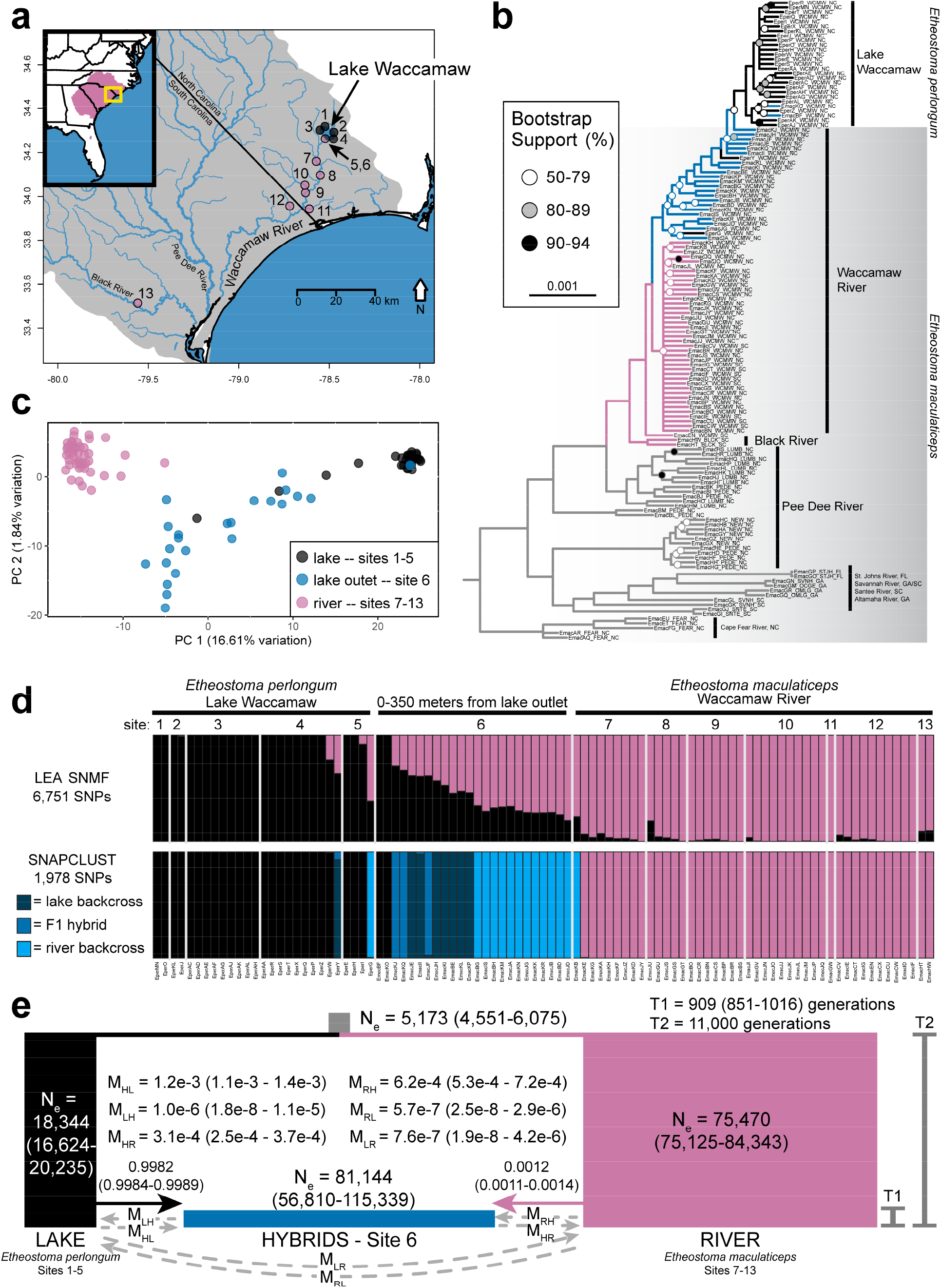
Evolutionary history of lake-river divergence. A) Map of sampling localities within the Waccamaw and Pee Dee River systems. Site 5 is located within Lake Waccamaw immediately above the outlet, site 6 is 0-350 meters downstream of the lake. Insert map shows range of *Etheostoma maculaticeps* in pink with sampling area highlighted in yellow. Gray areas indicate the Pee Dee River basin. B) Maximum likelihood ddRAD phylogeny of *E. maculaticeps* and *E. perlongum*. Outgroup taxa are not shown. Nodes with <50% bootstrap support are collapsed, nodes with >95% bootstrap support are unlabeled. Branches colored according to hybrid classification results. C), PCA of 10,628 SNPs with <10% missing data. D) Genetic clustering analyses. Each vertical bar represents a sample, bar colors represent admixture coefficients (SNMF) or assignment probabilities (SNAPCLUST). Sampling sites are delimited by vertical white bars. E) Parameter estimates from model with the best AIC score (Fig. S4, model 2, 11,000 generations of lake-river divergence). Non-parametric bootstrap 95% HDIs are indicated in parentheses.

### A *de novo* Darter Genome

We sequenced and assembled a *de novo* genome for *Etheostoma perlongum*. One *E. perlongum* individual (YFTC 37806, YPM ICH 033096) was euthanized with MS-222 and immediately dissected. We isolated fin, muscle, gill, liver, brain, eye, heart, gut, and testes tissue. Tissues were immediately frozen with liquid nitrogen and stored at −80 °C. For genome sequencing, we extracted DNA from muscle and gill tissue using a QIAGEN Genomic-tip 500/G kit. We used Oxford Nanopore long-read technology to sequence the *E. perlongum* genome. Prior to sequencing, we performed a size selection using the Circulomics Short Read Eliminator Kit. Sequencing libraries were prepared using 1 µg of gill or muscle genomic DNA and the Oxford Nanopore LSK-109 ligation sequencing kit. We also performed short-read Illumina sequencing with a 10x Genomics Chromium library preparation, allowing us to polish error-prone Nanopore reads with accurate short reads. The Nanopore data were assembled into contigs and scaffolded using Hi-C chromatin conformation capture sequencing (see Supporting Information for details).

To generate evidence for gene annotation, we performed transcriptome sequencing using eight different tissues (gill, fin, gut, heart, liver, brain, eye, and testes). We combined equimolar amounts of RNA from each tissue type for transcriptome sequencing performed by NovoGene (https://en.novogene.com/). We used the Maker v.2.3.10 pipeline to assign putative functional annotations to the *Etheostoma perlongum* genome assembly^26^. For annotation evidence, we used the transcriptome generated in this study, exon sequences from a close relative^27^ and protein sequence data from eight other teleost fishes. We performed four iterative rounds of annotation with Maker: one round of evidence-based annotation and three rounds *ab initio* gene prediction. Putative functions were assigned to the predicted genes by identifying potential matches to the UniProt Swiss-Prot database and searching for protein domains with InterProScan v.5^28^. The CoGe SynMap tool (https://genomevolution.org/coge/) was used to visualize synteny between the annotated *E. perlongum* genes two relatives in the family Percidae: *Etheostoma spectabile* and *Perca flavescens*. Details of the genome annotation pipeline are provided in the Supporting Information.

### Linkage Disequilibrium and Inversion Detection

Using the ddRAD assembly, we estimated linkage disequilibrium (LD) between all pairs of single nucleotide polymorphisms (SNPs) for the largest genomic scaffolds representing the 24 chromosomes. We created a reduced dataset using VCFTools v.0.1.15^29^, retaining only biallelic SNPs (“--max-alleles 2”), SNPs with less than 50% missing data (“--max-missing 0.5”), and SNPs with minor allele frequency > 5% (“--maf 0.05”). We estimated LD using PLINK v.1.90^30^, then averaged LD estimates in 1Mb windows across each scaffold for visualization. Additionally, following Matschiner et al. (2022), we calculated an individual SNP LD metric as the cumulative sum of distance (in base pairs) between other intrachromosomal SNPs in high LD (R^2^ > 0.8).

We used LDna v.2.0^31^ to identify clusters of ddRAD loci in high LD that could represent chromosomal rearrangements. We focused on “single outlier clusters” (SOCs) of high LD, as “compound outlier clusters” comprised of multiple SOCs can be the product of different evolutionary forces and are difficult to interpret^31^. LDna analyses were performed separately for the 24 largest scaffolds. To further examine SOC-1 on chromosome 9 (see Results), we recalculated LD as described above separately for samples with at least 75% of SOC-1 SNPs homozygous for the reference (lake) or alternate (river) allele. LDna details are provided in the Supporting Information.

### Phylogeny and Population Structure

To understand the broader evolutionary context of *Etheostoma perlongum*, we performed concatenated phylogenetic analyses of the ddRAD data using IQTree v.1.6.12^32^. To determine the impact of missing data on phylogenetic inference, we assembled alignments with 30%, 20%, 10%, and 5% missing data (Table S3). We used a GTR + gamma nucleotide substitution model for all analyses. We ran each analysis until 100 unsuccessful tree search iterations were completed. To assess topological support, we also performed 1,000 ultrafast bootstrap replicates^33^.

We assembled a set of filtered SNPs for 93 individuals from the Waccamaw River system plus two individuals from the Black River (a small tributary just near the mouth of the Waccamaw River) to assess population structure. Using VCFTools, we retained only biallelic SNPs (“--max-alleles 2”) and removed sites that contained more than 10% missing data (“--max-missing 0.9”). Since SNP singletons can confound inference of population structure^34^, we also removed SNPs with minor allele frequency < 5% (“--maf 0.05”). Finally, we thinned our dataset to include only a single SNP per 10,000 bp window (“--thin 10000”) to minimize the effects of physical linkage on our inference of population structure. We used PCA to examine general population structure patterns and sparse non-negative matrix factorization (sNMF) to estimate individual ancestry coefficients with the R package LEA v.2.6.0^35^. Based on the genetic clustering results from PCA and LEA, we used the *snapclust* function in the R package adegenet v.2.1.3^36, 37^ to classify individuals as hybrids or backcrosses. Additional details are available in the Supporting Information.

### Demographic Modeling

Demographic modeling using coalescent simulations was used to compare different evolutionary scenarios in the Waccamaw River basin. We used fastsimcoal2 v.2.6.0.3^38^ to estimate parameters from the observed site frequency spectrum (SFS). We compared three evolutionary models (Fig. S4A). All models contained three populations: *Etheostoma perlongum* in Lake Waccamaw, the hybrid population at the Lake Waccamaw outlet, and *Etheostoma maculaticeps* in the Waccamaw River. In each model, the hybrid population is formed by an admixture event after initial divergence between Lake Waccamaw and the Waccamaw River. The models varied in the number of migration edges connecting each population. In model one, no gene flow was allowed between the three populations (Fig. S4A). Model two allows for migration only after the formation of the hybrid zone, a scenario of secondary contact following allopatric divergence (Fig. S4A). Finally, model three allows continuous migration following initial divergence between Lake Waccamaw and the Waccamaw River, a scenario of parapatric divergence with gene flow (Fig. S4A). We compared model fit by calculating AIC scores. For the best-fit model, we generated 100 non-parametric bootstrap replicates and calculated the 95% highest density interval for each parameter. Details of model specification are provided in the Supporting Information.

### Genetic Geographic Clines

Genetic clustering analyses reveal broad population patterns, but different regions of the genome may have experienced different levels of gene flow between *E. perlongum* and *E. maculaticeps*. Therefore, we examined geographic clines in allele frequencies for 10,480 SNPs. We used the same VCFTools filtering parameters as for the population structure analyses but we did not thin SNPs by genomic position. Allele frequency clines were fit using the R package HZAR v.0.2.5^39^. For each SNP, we compared a null model (no difference in allele frequency between the lake and river) and three geographic cline models: a logistic curve with empirical maximum and minimum allele frequencies, a logistic curve with estimated maximum and minimum allele frequencies, and a logistic curve with estimated maximum and minimum allele frequencies and exponential decay curves at both ends. We compared these models using AICc scores. Model details are provided in the Supporting Information.

### Genetic Outlier Detection

Genetic loci under strong selection or with reduced gene flow during speciation may manifest as strongly differentiated outliers relative to the genomic background. We examined the ddRAD data for the presence of outlier loci using several different approaches. First, we assembled a SNP dataset that excluded individuals from site 6, since this locality consisted almost entirely of admixed hybrid individuals. The remaining individuals were classified as either lake (sites 1-5, n=27) or river (sites 7-13, n=44). We used VCFTools to retain only biallelic SNPs (“--max-alleles 2”), remove sites that contained more than 10% missing data (“-- max-missing 0.9), and remove SNPs with minor allele frequency < 5% (“--maf 0.05”). After filtering, the dataset contained 12,214 SNPs. We identified SNP outliers using a PCA-based approach^40^, and two methods utilizing the fixation index, F_ST_^41–43^. Since variation in sequencing depth can lead to imprecise F_ST_ estimates for individual SNPs^44, 45^, we also estimated relative divergence (F_ST_), absolute divergence (D_XY_), and nucleotide diversity (π) in 1 Mb sliding windows with a slide size of 100 Kb. Regions above the 99^th^ percentile of differentiation were designated as outliers. See Supporting Information for details.

### Gene Ontology Enrichment Analysis

To determine whether any gene ontologies (GO) were overrepresented regions of elevated divergence, we performed GO enrichment analysis in three genomic windows (F_ST_ plateau, D_XY_ peak 1, and D_XY_ peak 2, Fig. 4A). First, we assembled a reference database of amino acid sequences from three closely related species: *Larimichthys crocea* (Ensembl L_crocea_2.0), *Gasterosteus aculeatus* (Ensembl BROAD S1), and *Cottoperca gobio* (Ensembl fCotGob3.1). We used BlastX (NCBI) to perform a search against this reference database using the *Etheostoma perlongum* predicted transcriptome as the query sequence. We then used the GOATOOLS *find_enrichment.py* script to identify GO terms that were overrepresented in genes found within outlier regions^46^. REVIGO^47^ was used to cluster redundant GO terms with 50% allowed SimRel similarity. Additional details in the Supporting Information.

### Morphology

We used geometric morphometrics to assess the degree of body shape divergence between populations from Lake Waccamaw and the Waccamaw River. In total, we digitized a series of 16 landmarks and 24 sliding semi-landmarks (Fig. 2A) for 240 specimens using tpsDIG v.2.31^48^. We performed principal components analysis of the landmark data using the *PCA* function in the R package Momocs v.1.2.9^49^ and visualized morphospace with the Momocs *plot.PCA* function. In addition to the landmark analysis, we took four linear morphological measurements related to feeding ecology, swimming performance, and predator avoidance^50, 51^: head length, lower jaw length, body depth, and caudal length (Fig. 2A). We analyzed geographic variation in the second principal component axis (Fig. 3B), body depth, caudal length, head length, and upper jaw length (Fig. 2A) using a cline-fitting approach in the R package HZAR. The last four traits were corrected for body size by taking the residual versus standard length (Fig. 2A). We performed cline fitting as described for the SNP data.

**Figure 2.**
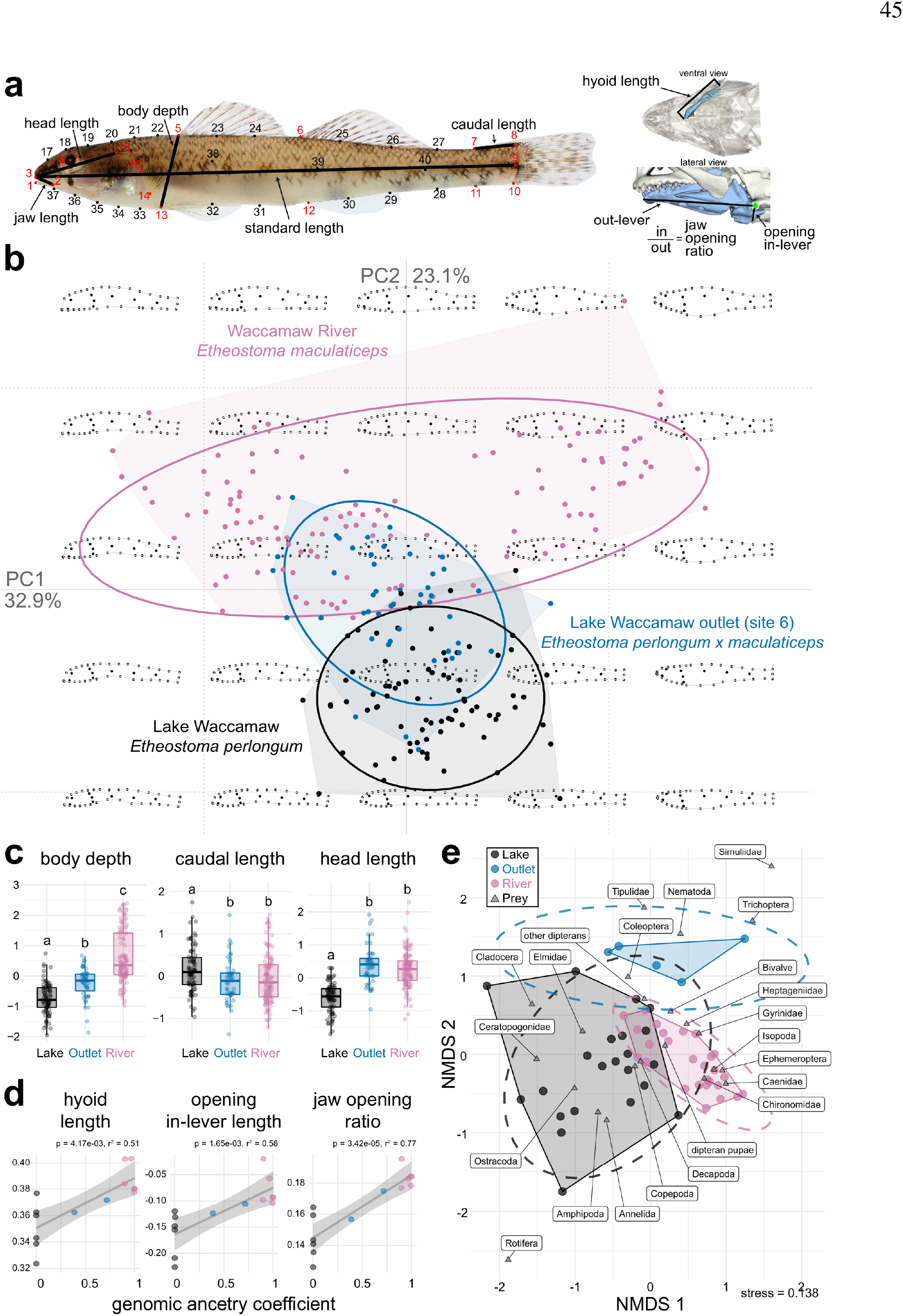
Morphological and ecological divergence associated with genomic divergence. A) Position of 16 landmarks (red) and 24 sliding semilandmarks (black) and the linear measurements. B) Morphospace of first and second principal components of the geometric morphometric analysis. Filled regions are convex hulls with 95% credible interval ellipsoids around the mean for each group. Gray outlines represent body shapes in different regions of morphospace. C) Boxplots of PC2 scores plus linear regression residuals of caudal length and head length versus standard length. Boxplots center line shows the median; box limits show 25th and 75th quartiles; whiskers show 1.5x interquartile range; all data points shown. Compact letter displays of Tukey HSD tests indicated above each plot. D) Linear regressions of osteological traits versus genomic ancestry coefficients estimated by SNMF (Fig. 1D). Points are colored by sampling location (lake, outlet, or river). E) Results of non-metric multi-dimensional scaling (NMDS) with fish stomach contents. Circles indicate fish specimens (colored by sampling location) while triangles represent prey categories. Filled regions are convex hulls with 95% credible interval ellipsoids around the mean for each group.

**Figure 3.**
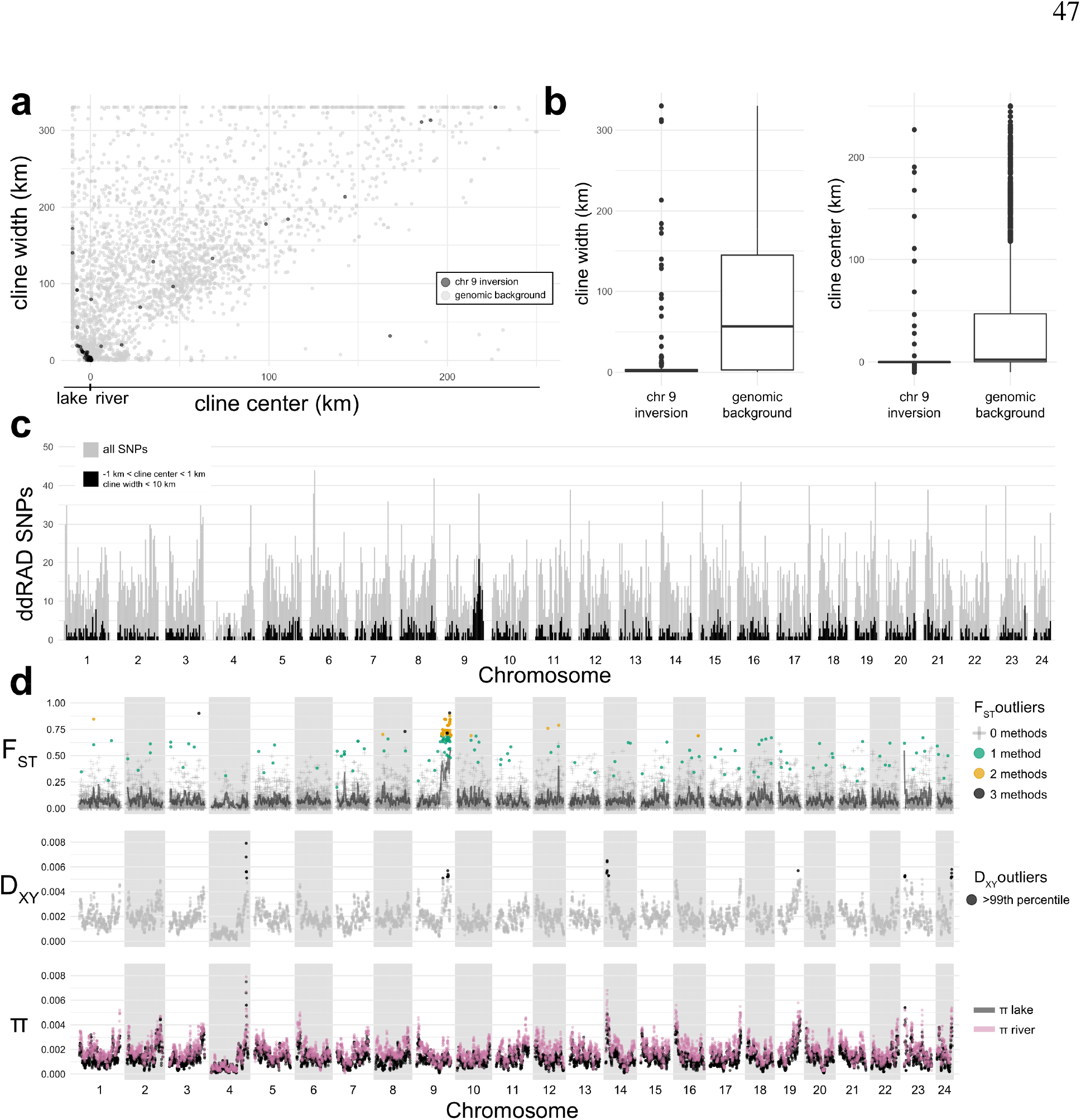
Genomic outliers of lake-river divergence are concentrated on chromosome 9. A) Scatter plot of estimated cline centers vs cline widths for each ddRAD SNP. The outlet of Lake Waccamaw is located at 0 km on the x-axis. Black points represent SNPs within the inversion. B) Boxplots of cline center and cline width for SNPs within the chromosome 9 inversion versus the genomic background. C) SNP density along the 24 largest *Etheostoma perlongum* scaffolds, histograms bins in 1 Mbp windows. Gray bars show the distribution of all analyzed SNPs. Black bars show the distribution of SNPs with estimated cline centers within 1 km of the lake-river boundary and with estimated cline widths < 10 km. D) Genome-wide estimates of F_ST_ (relative divergence), Dxy (absolute divergence), and π (nucleotide diversity) between Lake Waccamaw (*E. perlongum*) and the Waccamaw River (*E. maculaticeps*). F_ST_ estimates for individual SNPs are indicated by dots or crosses, sliding window F_ST_ estimates are indicated by the black line. Point color indicates how many methods identified a particular SNP as an outlier. Dxy plot shows sliding window estimates, black points indicate outliers. π plot shows sliding window estimates for Lake Waccamaw (black) and the Waccamaw River (pink). All sliding window estimates used a 1 Mbp window size with a 100 Kbp step size.

To further investigate ecomorphological patterns among river, lake, and hybrid-zone populations, we made linear measurements of the skeleton from µCT scans of a subset of 18 individuals (eight lake, six river, and four from the hybrid-zone). We made ten measurements that have been previously used to study darter comparative morphology^52^ and are expected to differ with changes in feeding ecology and diet. We also use our measurements to calculate two ratios, including jaw opening-lever ratio. Finally, we collected a series of meristic trait measurements that are traditionally used to discover and delimit species in ichthyology (Dataset S1). We compared morphometric, osteological, and meristic trait distributions among the lake, lake outlet, and river populations using ANOVA. For meristic and osteological traits, we also examined the correlation with lake-river genomic ancestry estimated from the SNP dataset. Additional details of measurements and analyses are found in the Supporting Information.

### Diet

We assessed the trophic niche of Lake Waccamaw and Waccamaw River populations by collecting diet data from gut contents of specimens of *E. perlongum* (n=22), the hybrid zone (n=5), and *E. maculaticeps* (n=27). Contents of the foregut were identified to ordinal or family level, where possible. We estimated multivariate trophic ecospace using Nonmetric Multidimensional Scaling (NMDS) and visualized the distribution of prey items with a bipartite network. Additional details are provided in the Supporting Information.

## RESULTS

### Genome Assembly

We produced a de novo chromosome-level genome assembly of *Etheostoma perlongum* using Nanopore long reads polished with high-quality Illumina short reads and scaffolded with a Hi-C contact map (Table S3). We generated 29.85 Gb of Nanopore sequence data, with a median read length of 2,634 bp and a read length N50 of 10,459 bp. 1% of the Nanopore reads are longer than 100 Kbp, with a maximum length of 301,807 bp. Based on the genome assembly size of another *Etheostoma* species^53^, our Nanopore sequence data represents approximately 35X genomic coverage for *E. perlongum*. The total length of the 150 bp Illumina paired-end reads used for error correction is 111.26 Gb, representing an additional approximately 130X genomic coverage.

The total size of the scaffolded *E. perlongum* genome assembly is 788 Mb, slightly smaller than other assembled percid genomes (958 Mb, *Perca fluviatilis*^54^; 877 Mb, *Perca flavescens*^55^; 855 Mb, *Etheostoma spectabile*^53^). The *E. perlongum* genome assembly is comprised of 2,095 contigs (N50 = 2.3 Mb, L50 = 100 contigs) grouped into 1,179 scaffolds (N50 = 30.8 Mb, L50 = 11 scaffolds). The 24 longest scaffolds account for 92.4% of the total assembly length and exhibit a 1:1 correspondence with *Perca flavescens* and *Etheostoma spectabile* chromosomes (Fig. S1). Hereafter, we refer to the 24 longest *E. perlongum* scaffolds as chromosomes. The *E. perlongum* genome assembly exhibits conserved synteny with the *Perca flavescens* genome^55^ and, to a lesser extent, the *Etheostoma spectabile* genome^53^ (Fig. S1).

96.7% of BUSCO Actinopterygii orthologs are represented as complete sequences in the *E. perlongum* genome assembly, with an additional 1% represented as partial sequences. The annotation pipeline identifies 42% (328.6 Mb) of the *E. perlongum* assembly as repetitive DNA, a larger proportion than the 30-33% of repetitive DNA identified in other percid genomes^53–55^. Total GC content of the *E. perlongum* genome is 40.5%. The *E. perlongum* transcriptome consists of 260,143 transcripts for 138,096 genes, with a contig N50 of 1,731 bp. 71.5% of BUSCO Actinopterygii orthologs are represented as complete sequences in the transcriptome, with an additional 12.2% represented as partial sequences. Using the *E. perlongum* transcriptome and several other lines of evidence, *ab initio* genome annotation with the Maker pipeline identifies 28,034 putative protein-coding genes. 80% (22,534) of these predicted genes are assigned putative functional annotation based on homology with the UniProt Swiss-Prot database. 75% (16,850) of matches with the UniProt Swiss-Prot database are longer than 30 amino acids and have >50% sequence similarity. Additionally, InterProScan identifies 41,636 functional domains for 19,948 of the predicted genes.

### Phylogenetics and Population Genetics

We used the *E. perlongum* genome to map and assemble ddRAD sequence data from 140 individuals of *E. perlongum* and *E. maculaticeps*. All individuals in the ddRAD dataset are represented by at least 461,000 reads, with a substantially larger mean number of reads per sample (mean = 3.7E+6, sd = 2.1E+6). The assembly pipeline identifies 267,237 orthologous ddRAD loci shared by at least four samples. In the filtered dataset containing only biallelic SNPs with minor allele frequency greater than 5% and fewer than 10% missing genotypes, 98% (10,628/10,752) of the SNPs are located on the 24 longest genomic scaffolds.

Despite considerable variation in the percent of missing data (Table S4), analyses of all alignments produce well-resolved phylogenies with congruent relationships between populations of *Etheostoma perlongum* and *E. maculaticeps.* We therefore report only results from the phylogenetic analysis of the dataset with the largest number of ddRAD loci. All individuals from the Waccamaw River and Lake Waccamaw resolve in a strongly supported clade that is nested within a lineage comprised of populations from the lower Pee Dee River system (Fig. 1B).

However, genetic differentiation exists at fine spatial scales between Lake Waccamaw and the Waccamaw River (Fig. 1C,D). Within the Waccamaw basin, the first principal component (PC) axis of genetic variation separates individuals from Lake Waccamaw and the Waccamaw River. Likewise, cross-entropy scores estimated by sNMF exhibit an “elbow” at K=2, indicating that two ancestry clusters best explain the genetic data (Fig. S2A). Analyses with K>2 do not reveal additional population substructure within the Waccamaw basin (Fig. S2B). In addition to the two genetic ancestry clusters, we find evidence of an active hybrid zone in the river outlet immediately downstream of Lake Waccamaw (site 6, Fig. 1A). Most individuals at this site have intermediate PC1 scores (Fig. 1C) and exhibit admixed ancestry (Fig. 1D). The snapclust algorithm classifies all but two individuals from this site as putative hybrids or backcrosses (Fig. 1D). Outside of locality 6, only three other individuals are classified as hybrids or backcrosses.

### Demographic Modeling

To date the origin of *Etheostoma perlongum* and examine the history of gene flow between *E. perlongum* and *E. maculaticeps*, we compared three different demographic scenarios across a range of plausible divergence times (Fig. S3a). The first scenario models allopatric divergence with a single instance of secondary contact to form a hybrid zone, but no subsequent gene flow. The second scenario also models allopatric divergence with secondary contact but allows gene flow between the two species after the formation of a hybrid zone. The third scenario models parapatric divergence by allowing continuous gene flow between the two lineages immediately after their initial split.

The scenarios of secondary contact with gene flow and parapatric divergence have consistently better Akaike information criterion (AIC) scores than the model with no gene flow after secondary contact (Fig. S3b). Support for these two models is mixed across a range of initial divergence times (ΔAIC < 10, Fig. S3c, Table S5). The most strongly supported demographic models indicate that divergence between *Etheosotma perlongum* in Lake Waccamaw and *E. maculaticeps* in the Waccamaw River began 11,000 to 15,000 generations ago (Table S5). With an Akaike weight of 0.89, the best fit of all examined models is the scenario of allopatric divergence between *E. perlongum* and *E. maculaticeps* 11,000 generations ago with secondary contact and gene flow in the last 909 generations (Fig. 2E).

Demographic inference suggests that a spatially narrow hybrid zone facilitates gene flow between *Etheostoma perlongum* and *E. maculaticeps* (Fig. 2E). Under the best fit model, direct migration rate estimates between *E. perlongum* and *E. maculaticeps* are very low (lake to river N_e_m [expected number of migrants per generation] ∼= 0.06, river to lake N_e_m ∼= 0.01). However, there are high migration rates from *E. maculaticeps* in the Waccamaw River into the hybrid population (N_e_m ∼= 50), as well as a slightly lower migration rate in the reverse direction (N_e_m ∼= 23). The opposite is true for E. *perlongum* in Lake Waccamaw, with a migration rate from the hybrid population into *E. perlongum* (N_e_m ∼= 22) that is much higher than the migration rate from *E. perlongum* into the hybrid population (N_e_m ∼= 0.08).

### Ecological and morphological divergence

We examined several lines of evidence for lake-river morphological and ecological divergence. Geometric morphometric analyses of body shape reveal that the first principal component (PC) axis largely corresponds to a concave or convex shape of the abdominal region, which can be explained by the presence of gravid females in the dataset (Fig. S4). However, PC2 differentiates Lake Waccamaw *Etheostoma perlongum* from Waccamaw River *E. maculaticeps* (Fig. 2B). Lake individuals have more slender bodies and elongate caudal regions compared to individuals from the river (Fig. 2B). There is also an associated change in mouth position and head shape, with lake fish having more terminal mouths than their river counterparts (Fig. 2B). Individuals from the hybrid zone downstream of Lake Waccamaw exhibit intermediate body shapes (Fig. 2B). Body depth contributes strongly to this intermediacy; individuals from the hybrid zone are distinct from both *E. perlongum* in Lake Waccamaw and the Waccamaw River population of *E. maculaticeps* (Fig. 2C). Body depth, head length, and PC axis two exhibit sharp geographic clines centered at the lake-river boundary (Fig. S5).

µCT imaging reveals that *E. perlongum* and *E. maculaticeps* differ in the number of vertebrae and several ecologically relevant measures such as the length of the hyoid and jaw opening in-lever (Fig. 2D, Fig. S6). In addition, meristic traits that are traditional ichthyological taxonomic characters differ between *E. perlongum* and populations of *E. maculaticeps* in the Waccamaw River (Fig. S8). Several osteological and meristic traits that differ between the lake and river are strongly correlated with genomic ancestry estimates, and admixed individuals have intermediate trait values (Fig. 2D, Fig. S6,7).

To further explore ecological lake-river divergence, we examined the gut contents of *E. perlongum* and Waccamaw River *E. maculaticeps*. We identified the gut contents from 25 specimens of *Etheostoma perlongum*. Two of these specimens were gravid females with empty stomachs and were excluded from further analysis. The three most frequent prey items to occur in *E. perlongum* stomachs are Amphipods, Chironomidae midge larvae, and Ostracods, each appearing in more than 50% of examined stomachs. Copepods, Cladocera, and Ceratopogonidae larvae also occur in greater than 10% of stomachs examined. A variety of benthic invertebrates occur in two or fewer individual stomachs, including Ephemeroptera larvae, Elmidae larvae, Diptera pupae, Annellidae, Rotifera, and a Decapoda (Dataset S3).

We identified contents of 28 stomachs from *E. maculaticeps* specimens (one empty and excluded from analysis), and six specimens from the contact zone in the Waccamaw River, just below Lake Waccamaw (one with empty stomach and excluded from analysis). Chironomidae larvae are present in all *E. maculaticeps* stomachs, and Isopods (52%) and Ephemeroptera larvae (48%) are the other most frequently consumed prey. Trichoptera and Coleoptera larvae, Cladocera, Copepods, Amphipods, and Ostracods are consumed by 10-20% of riverine individuals.

*Etheostoma perlongum* and *E. maculaticeps* consume many of the same prey categories, but their strength of association with these common prey differed (Dataset S3, Fig. S9). The percent frequency occurrence of Chironomidae larvae is high for both *E. perlongum* and *E. maculaticeps*, but stomachs of *E. maculaticeps* tend to contain more chironomid larvae (mean 16.37, maximum 44) than *E. perlongum* (mean of 5.19, maximum 18). Microcrustacea (Amphipods, Ostracods, Copepods) are consumed by both species, but occur in *E. perlongum* stomachs more frequently and in higher numbers. Bipartite network analysis associates Ephemeroptera, Trichoptera, Coleoptera, Chironomidae, Bivalves, and Isopoda with *E. maculaticeps*, while the other prey categories (e.g., Ceratopogonidae, Amphipoda, Ostracoda, Copepoda) are more strongly associated with *E. perlongum* (Fig. S9).

### Outlier SNPs

We examined thousands of SNPs to identify regions of the genome that are resistant to gene flow between *Etheostoma maculaticeps* and *E. perlongum*. Geographic cline analyses reveal that 14.6% of the 10,752 SNPs are best described by a model with a sharp allele frequency cline less than 10 km wide and centered within 1 km of the lake-river boundary (Fig. 3A). While SNPs are broadly distributed across the genome, SNPs with the narrowest clines centered on the lake-river boundary are concentrated on chromosome 9 (Fig. 3B,C). 33% of the SNPs on chromosome 9 exhibit narrow geographic clines centered within 1 km of the lake-river boundary, while no other chromosome has more than 20% of SNPs with such narrow clines. This pattern is most pronounced within the last 9 Mb of chromosome 9, where 57% (110 of 195) of SNPs exhibit clines narrower than 10 km centered within 1 km of the lake-river boundary (Fig. 3A,C).

Although SNPs with high lake-river differentiation are scattered across the genome, outlier detection methods reveal an overabundance of highly differentiated SNPs within the last 9 Mb of chromosome 9 (Fig. 3D). 45% (86 of 193) of PCAdapt outliers, 85% (53 of 62) of SNP F_ST_ outliers, and 67% (4 of 6) of BayeScan outliers are found on chromosome 9. The abundance of high F_ST_ SNPs on chromosome 9 is not the result of biased sex-ratio sampling; separate analyses of males and females qualitatively reveal the same patterns (Fig. S9). Outlier SNPs are not evenly distributed across chromosome 9; 94% (81 of 86) of chromosome 9 outlier SNPs occur within a 9 Mb plateau of elevated divergence with no obvious peak (Fig. 4A). Sliding window estimates reveal a similar pattern: 86% (31 of 36) of all outlier F_ST_ windows create a plateau of divergence along the last 9 Mb of chromosome 9 (Fig. 4A). Within the 9 Mb plateau of high F_ST_, there are two smaller outlier peaks of elevated D_XY_ (Fig. 4A). The 9 Mb region of chromosome 9 is the only genomic region that exhibits both elevated relative (F_ST_) and absolute (D_XY_) lake-river divergence (Fig. 3D).

**Figure 4.**
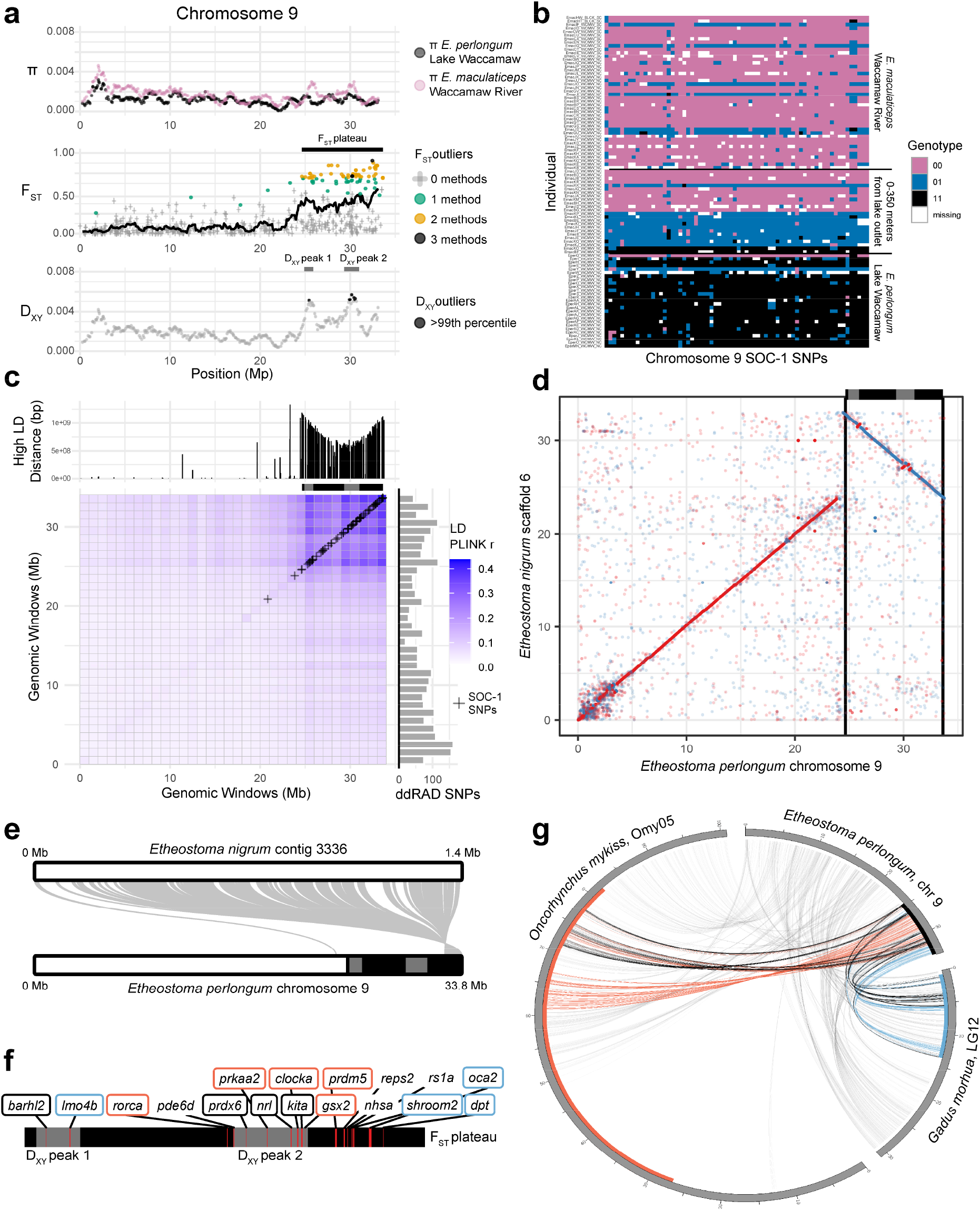
*Etheostoma perlongum* chromosome 9 contains a likely inversion supergene. A) Chromosome 9 patterns of nucleotide diversity (π) and genomic differentiation (F_ST_ and Dxy). F_ST_ estimates for individual SNPs are indicated by dots or crosses, sliding window F_ST_ estimates are indicated by the black line. Point color indicates how many methods identified a particular SNP as an outlier. Dxy and π plots show sliding window estimates with a 1 Mbp window size with a 100 Kbp step size. The F_ST_ plateau (black bar) and Dxy peaks (grey bars) indicate outlier regions used for gene ontology enrichment analyses. B) Genotypes of SNPs in the chromosome 9 single outlier cluster of high LD (SOC-1). Individuals are ordered by increasing downstream distance from the north shore of Lake Waccamaw. C) Linkage disequilibrium (LD) heatmap for 1 Mbp windows along chromosome 9. Locations of SNPs in SOC-1 are noted. Bar plot on the top margin shows per-SNP linkage metric calculated as the sum of distances between SNPs in high LD (R^2^ > 0.8). Histogram on the right margin shows the distribution of ddRAD SNPs in 1 Mbp windows along chromosome 9. D) Nucmer alignment dot plot of *E. perlongum* chromosome 9 versus *E. nigrum* scaffold 6. Red indicates forward strand alignments, blue indicates reverse strand alignments. E) Alignment of *E. nigrum* contig 3336 spanning the *E. perlongum* inversion. F) Position of vision, olfaction, and circadian genes within the chromosome 9 F_ST_ plateau. G) Circos plot of gene synteny between *Etheostoma perlongum* chromosome 9, *Gadus morhua* linkage group 12, and *Oncorhynchus mykiss* chromosome 5. Red, blue, and black regions on each chromosome represent inversion/supergene location. Each link represents a syntenic gene pair. Gray links are genes that are not syntenic between two inversions. Orange links indicate genes syntenic between the *E. perlongum* and *O. mykiss* inversions, blue links indicate genes syntenic between the *E. perlongum* and *G. morhua* inversions, and black links indicate genes syntenic between *E. perlongum* and both *O. mykiss* and *G. morhua* inversions.

### Evidence for an Inversion on Chromosome 9

To look for signals of structural genomic variation, we examined linkage disequilibrium (LD) using several SNP datasets. The 9 Mb high F_ST_ plateau on chromosome 9 has elevated LD compared to the genomic background (Fig. S10, Fig. 4C) and SNPs in this region comprise a single outlier cluster of high LD (SOC-1, Fig. 4C). SOC-1 SNPs are in very high LD (median intracluster LD = 0.88) and exhibit genotypes strongly associated with the lake and river lineages (Fig. 4B). Indeed, when lake and river chromosome 9 genotypes are examined separately, the 9 Mb block of high LD is no longer detected (Fig. S11, S12). The per-SNP LD metric shows a clear drop in linkage just beyond the boundaries of the F_ST_ plateau (Fig. 4C). This signature is also observed on chromosome 8 (Fig. S10), which also contains the only other outlier clusters of high LD (Fig. S13). However, unlike on chromosome 9, the high LD region on chromosome 8 persists when examining only river genotype individuals (Fig. S12) and is not strongly associated with divergent lake and river genotypes (Fig. S13).

We aligned the *E. perlongum* genome to a de novo, chromosome-level assembly of a close relative, *E. nigrum.* Compared to *E. nigrum* scaffold 6, there is a clear inversion at the end of *E. perlongum* chromosome 9 (Fig. 4D). This large inversion corresponds precisely with the 9 Mb high F_ST_ plateau. The inversion breakpoint on *E. perlongum* chromosome 9 occurs at approximately 23.84 Mb and is spanned by a single *E. nigrum* contig (Fig. 4E). Unlike the chromosome 9 inversion, the region of high LD on *E. perlongum* chromosome 8 is not inverted relative to the *E. nigrum* genome (Fig. S1).

Annotation of the *E. perlongum* genome assembly identifies key genes linked to vision and olfaction within the chromosome 9 inversion, many of which are concentrated within the two D_XY_ peaks (Fig. 4F, Table S6). Several of these genes, such as *pde6d*, *prdx6,* and *prdm5*, are related to severe retinal disease phenotypes in vertebrates^56–58^ (Table S6). Gene ontology (GO) enrichment analyses also identify many biological processes that may be overrepresented within the *E. perlongum* chromosome 9 inversion (Table S7), though controlling for FDR results in no significant enrichment. Strikingly, the *E. perlongum* chromosome 9 inversion contains many genes, including key vision genes, that are also present in a well-document inversion supergene on Atlantic Cod linkage group 12 and a massive double inversion supergene on Rainbow Trout chromosome 5 (Fig. 4G, Table S6, Dataset S4)^59, 60^.

## DISCUSSION

### Rapid Lacustrine Speciation with Gene Flow

Shallow genetic differentiation is unusual among closely related darter species, which are typically millions of years divergent and exhibit reciprocal monophyly^19^. However, unlike the geologically ancient rivers of southeastern North America home to most darter species, Lake Waccamaw was completely inundated during high sea levels in mid-Pliocene (2.9-3.3 Ma)^61^. The present configuration of Lake Waccamaw is relatively young; studies of sediment cores and pollen records estimate the lake originated in the late Pleistocene, 15,000-32,000 years ago^14, 62^. Under the best fit demographic model with a generation time of one to three years^63, 64^, speciation of *E. perlongum* initiated in the late Pleistocene or early Holocene (11,000 to 33,000 years ago), remarkably consistent with the young geologic age of Lake Waccamaw.

Although there is a narrow, contemporary hybrid zone between *E. perlongum* and *E. maculaticeps*, demographic analyses support an initial period of divergence with no gene flow (Fig. S3). Drought records show that Lake Waccamaw was occasionally isolated from the Waccamaw River^66^. Historically, the outlet of Lake Waccamaw may have been smaller and intermittent like the outlets of other Carolina Bay lakes^11, 12^. We hypothesize that historic fluctuations in connectivity between the lake and river created periodic microgeographic isolation, limiting gene flow and initiating speciation.

Speciation frequently does not lead to complete reproductive isolation^65^. In darters, hybridization and introgression are common^19^; approximately 25% of darter species are involved with natural hybrid crosses^67^. However, hybridization at an early stage of speciation is unusual, as 93% of darter sister species pairs are allopatric, precluding opportunities for gene flow^19^. Even recently diverged darter species such as *Percina freemanorum* and *P. kusha* (diverged approximately 300,000 years ago) are allopatric and do not actively hybridize^68^. Geographic isolation is the main driver of darter diversification^19, 69^ and most darter hybridization follows long periods divergence^70^. For example, hybridization between *E. caereuleum* and *E. spectabile* is well-characterized on a phenotypic and genomic level^53^, yet these species share most recent common ancestry 22 million years ago^19^. In contrast, *E. perlongum* and *E. maculaticeps* diverged within the past 8,000 to 39,000 years (Fig. 1D) but are actively hybridizing. Thus, this young species pair is the only known example among darters where gene flow can occur early in the speciation process. However, demographic analyses and geologic evidence indicate that microgeographic isolation still played an important role during the onset of speciation.

### Ecology and Morphology Reflect Genetic Divergence

Morphological and ecological divergence provide additional evidence that *Etheostoma perlongum* is a distinct species. *Etheostoma perlongum* differs from *E. maculaticeps* along multiple ecomorphological axes, including an elongated and shallower body (Fig. 2). In fishes, slender body shapes are often associated with sustained swimming speeds, whereas deeper bodies can allow for more maneuverability in complex habitats^50^. Lake Waccamaw is an open lacustrine environment where sustained swimming performance may facilitate predator avoidance^71^. Morphologically and genetically intermediate hybrid individuals in the lake outlet mark the center of a steep, narrow geographic cline (Fig. S5), which suggests divergent selection is acting on these phenotypes in lentic versus lotic habitats.

In addition to an elongated body shape, differences in head and jaw shape between *E. perlongum* and *E. maculaticeps* indicate divergence in foraging ecologies. Lakes typically have an abundance of limnetic prey, while rivers offer mostly benthic prey; these differences in prey availability drive lake-river body shape divergence in other fish species^72^. Analysis of gut contents reveals that the two species occupy distinct trophic niches (Fig. 2E), with *E. perlongum* associated more strongly with microcrustaceans from the water column such as amphipods, ostracods, and copepods^73^. These prey taxa appear infrequently in the diets of *E. maculaticeps*, which has a stronger association with strictly benthic prey taxa^73^ that rarely (Ephemeroptera) or never (Trichoptera and Isopoda) occur in the diets of *E. perlongum* (Fig. S8).

The combination of a more terminal mouth, an elongated body, and stronger association with limnetic prey (Fig. 2) suggests that *Etheostoma perlongum* has partially escaped the benthic lifestyle typical of most darter species and utilizes more prey items in water column. Moreover, *E. perlongum* has a smaller lever ratio for lower jaw opening than *E. maculaticeps* (Fig. 2D). This indicates enhanced mobility of the lower jaw associated with suction feeding compared to high values which are force-enhanced and associated with biting^74^. Other darter species associated with sandy microhabitats like Lake Waccamaw also have smaller jaw opening-lever ratios, which may aid in suctioning prey from loose substrate or feeding on more evasive zooplankton^52^. In addition to ecological and morphological divergence, *Etheostoma perlongum* exhibits an annual life cycle and faster growth rates compared to a generation time of 2 to 3 years in typical riverine populations of *E. maculaticeps*^63, 64^. We hypothesize that the annual life history of *E. perlongum* may be related to greater seasonality of prey or differential predation pressures in lacustrine versus riverine ecosystems^75^.

### Divergence Concentrated in a Large Chromosomal Inversion

Divergence across the genome is often heterogeneous during speciation^76^ and many models of ecological speciation invoke genomic “islands” or “continents” of differentiation^77^. We observe a single 9 Mb “continent” of genomic divergence between *E. perlongum* in Lake Waccamaw and *E. maculaticeps* in the Waccamaw River (Fig. 3, 4) that appears resistant to otherwise high levels of lake-river gene flow (Fig. 1). This region contains a plateau of SNPs with high lake-river F_ST_ (Fig. 3D, 4A) and extremely steep and narrow geographic allele frequency clines centered at the lake-river boundary (Fig. 3A-C). Theory predicts that genomic regions resistant to gene flow should exhibit high levels of relative and absolute divergence^44^. We find that the only genomic windows with significantly elevated relative (F_ST_) and absolute (D_XY_) divergence occur inside the 9 Mb F_ST_ plateau on chromosome 9 (Fig. 3D). Within chromosome 9 and across the genome, we observe more F_ST_ outliers than D_XY_ outliers (Fig. 3D), matching theoretical predictions that relative divergence metrics are more responsive to reduced gene flow than absolute divergence metrics^44^. Importantly, the F_ST_ plateau of chromosome 9 does not exhibit reduced genetic diversity (Fig. 3D, 4A), which can inflate estimates of F_ST_^44^.

Why are collinear SNPs on chromosome 9 resistant to gene flow between *Etheostoma perlongum* and *E. maculaticeps*? One probable explanation is that a chromosomal inversion suppresses recombination in this region, maintaining tight linkage of *E. perlongum* and *E. maculaticeps* genotypes despite homogenizing gene flow. Models of inversion speciation have broad theoretical and empirical support^78, 79^. We demonstrate that the F_ST_ plateau on chromosome 9 contains an outlier cluster of high LD SNPs (Fig. 4C), with alternate genotypes segregating by lake or river habitat (Fig. 4B). Outlier clusters of high LD are linked to known chromosomal inversion in *Anopheles* mosquitos^31^, sticklebacks^80^, and *Littorina* snails^81^. The 9 Mb region of high LD on *Etheostoma perlongum* chromosome 9 spans several contigs and does not coincide with a scaffold breakpoint (Fig. S14), indicating this signature of high LD is not the result of genome assembly errors^82^.

Most connivingly, alignment of the *E. perlongum* genome to a close relative, *E. nigrum,* reveals a chromosomal inversion that aligns precisely with the high F_ST_ plateau (Fig. 4D, E). *Etheostoma nigrum, E. maculaticeps,* and *E. perlongum* share most recent common ancestry approximately 6 million years ago^83^. We hypothesize *E. nigrum* represents the ancestral arrangement of chromosome 9, and the 9 Mb inversion arose recently in *E. perlongum.* However, as chromosomal inversions are often polymorphic within species^59, 60^, future genome sequencing of *E. maculaticeps* is needed to confirm this hypothesis.

We propose that the large inversion on chromosome 9 suppresses recombination between Lake Waccamaw *E. perlongum* and Waccamaw River *E. maculaticeps* genotypes, providing a potential genomic mechanism to explain divergence and speciation despite gene flow. However, it is challenging to demonstrate that the chromosome 9 inversion played a direct role in speciation between *Etheostoma perlongum* and *E. maculaticeps*. Different mechanisms such as ancestral polymorphism can mislead inference about the importance of inversions in speciation^84^. Alternatively, proximity to the centromere could also produce high LD on chromosome 9 even without an inversion. Centromeric regions can play a role in speciation^85^, though sometimes incidental due to slow segregation of ancestral alleles^86^. However, if the 9 Mb region of chromosome 9 exhibits elevated LD simply because it is a centromeric region, we would expect to observe more regions of the *E. perlongum* genome with similarly high LD. Additionally, the 9 Mb region of chromosome 9 does not exhibit other hallmarks of centromeric regions, such as reduced gene density or increase density of repetitive sequences.

### Targets of Selection and Inversion Supergene Convergent Evolution

Genotypic and phenotypic divergence along narrow geographic clines is consistent with a scenario of divergent selection between *Etheostoma perlongum* in Lake Waccamaw and *E. maculaticeps* in the Waccamaw River^87^. In addition to predator avoidance and prey availability discussed above, researchers have long hypothesized that phenotypic divergence in the Waccamaw system is due to differences in light availability between the clearwater Lake Waccamaw and tannic or blackwater Waccamaw River^15, 18^. Water clarity and chemistry ecotones are a major driver of speciation in other fishes, especially in the Amazon basin^88^. Clearwater will favor greater use of vision for foraging, while blackwater favors olfactory development, particularly since the flowing water can transmit olfactory signals over longer distances than in standing water^89, 90^. The generality of lake-stream differentiation with respect to vision and olfaction is unclear, but there is some previous evidence of this tradeoff across similar environmental gradients^91^. Recent work demonstrates that the number of optic nerve fibers, rod, and cone cells differs between two other Lake Waccamaw endemic fish species, *Fundulus waccamensis* and *Menidia extensa*, and their riverine relatives^92^.

The *E. perlongum* chromosome 9 inversion contains numerous potential targets of divergent lake-river selection. Inversions suppress recombination, which can create supergenes of functionally linked loci that facilitate local adaptation in the face of gene flow^93^. Chromosomal inversions associated with local adaptation and ecological selection have been recently identified in several fish species^59, 60, 94, 95^. We hypothesize that the *Etheostoma perlongum* chromosome 9 inversion is a supergene containing numerous loci involved in retinal formation and olfaction that are likely targets of selection in lentic clearwater versus lotic blackwater habitats (Fig. 4E, Table S6). Additionally, the inversion contains genes involved in regulation of circadian rhythms such as *clocka*, which has been implicated in fish reproductive seasonality^96^ and may be linked to life history differences between the annual, semelparous *E. perlongum* and the perennial, iteroparous *E. maculaticeps*. Together, these linked genes are protected from homogenizing gene flow by an inversion and may have driven phenotypic and ecological divergence, facilitating speciation between *E. perlongum* and *E. maculaticeps*.

Surprisingly, the *E. perlongum* chromosome 9 inversion exhibits gene synteny with two other recently documented inversion supergenes in Atlantic Cod (*Gadus morhua*) and Rainbow Trout (*Oncorhynchus mykiss*)^59, 60^. Several genes, including *barlh2*, *prdx6*, *nrl*, and *kita* are present in the inversions of all three species. The similarity of gene content in these inversion supergenes is remarkable considering cod, darters, and trout share a most recent common ancestor in the middle Triassic, approximately 225 million years ago^97^. These regions also share deep homology with the mammalian X-chromosome, which contains many of the same retinal development and circadian rhythm genes. Divergence of ancestral mammalian X and Y chromosomes was also potentiated by an inversion, which disrupted recombination and allowed divergence through sex-antagonistic selection^98^. In fishes, the same molecular mechanism (chromosomal inversion) in a homologous region of the genome (a portion of the ancestral X-chromosome) has repeatedly facilitated divergence through population-antagonistic selection. In Atlantic Cod and Rainbow Trout, these supergenes are polymorphic across hundreds of kilometers and associated with local adaptation (cod) or divergent migratory behavior (trout)^59, 60^. In contrast, genotype frequencies within the *Etheostoma perlongum* chromosome 9 inversion shift drastically over the span of just a few kilometers (Fig. 3C, Fig. 4D). Thus, inversions in the same genomic regions containing similar sets of genes underlie both local adaptation across broad spatial scales and ecological speciation at fine spatial scales, demonstrating that convergent evolution of genomic architecture can lead to disparate outcomes.

### Conclusions

Together, our results uncover an atypical example of lacustrine speciation in darters. The *Etheostoma perlongum* genome assembly in combination with ddRAD sequencing reveals that *E. perlongum* in Lake Waccamaw is a young species that originated following the Pleistocene, with low levels of genetic differentiation from its sister lineage *E. maculaticeps* in the Waccamaw River. *Etheostoma perlongum* also exhibits subtle but significant differences in functional traits associated with body shape, the feeding system, and diet. Morphological and genetic intermediacy indicate an active hybrid zone between *E. perlongum* and *E. maculaticeps* in a short stretch of the Waccamaw River just downstream of Lake Waccamaw. While there is strong evidence of historical and contemporary gene flow, a 9 Mb genomic region on chromosome 9 appears resistant to introgression between *E. perlongum* and *E. maculaticeps*. This region contains an abundance of SNPs with steep, narrow allele frequency clines and elevated lake-river divergence relative to the genomic background. Elevated linkage disequilibrium in this region suggests that a chromosomal inversion is suppressing recombination, allowing alleles important for local adaptation or reproductive isolation to persist in the face of gene flow. This likely inversion may constitute a supergene involved in retinal development, olfaction, and circadian regulation, all potential targets of divergent lake-river selection. *Etheostoma perlongum* demonstrates how, even in clades dominated by allopatric diversification, structural genomic variation can overcome the effects of homogenizing gene flow and enable rapid ecological speciation.

## Acknowledgements

A. Dornburg, L. L. Bowman, A. Ghezelayagh, D. Kim, and M. Correa assisted in field collections. G. Watkins-Colwell secured collecting permits and provided museum collections support. B. Tracy (NC DEQ) and G. Hogue (NCSM) provided tissue samples for this work. The Yale Center for Genome Analysis and T. Lan assisted sequencing efforts for the *Etheostoma perlongum* genome. Lastly, the authors thank V. A. Albert, O. Gokcumen, and members of the Near, Donoghue, Muñoz, and Krabbenhoft labs for their feedback on earlier versions of the manuscript. This material is based upon work supported by the NSF Postdoctoral Research Fellowships in Biology Program under Grant No. 2109761.

