## Supporting Information for "Lacustrine speciation associated with chromosomal inversion in a lineage of riverine fishes"

### SI METHODS

#### ddRAD Sequencing

Genomic DNA was extracted from tissues using a standard DNAeasy Qiagen Blood and Tissue Kit. Prior to library preparation, we cleaned DNA extractions using an ethanol precipitation. 3M sodium acetate (pH = 5.2) was added equal to 10% of the total volume of the DNA extraction. We then added 100% ethanol equal to 2.5 times the total volume of DNA, mixed by inverting the tube, and froze for 10 minutes at -80 °C. Samples were centrifuged for 30 minutes at 8,000 RCF. We immediately poured off the supernatant and washed the DNA pellet with 250 uL of cold 70% ethanol. The tube was centrifuged again for 5 minutes at 8,000 RCF, then the supernatant was poured off. We allowed the pellet to air dry for about 15 minutes, then resuspended with the desired amount of DNAse-free water.

We performed a modified version of a double digest restriction site associated DNA (ddRAD) sequencing protocol[1,2]. We digested 200 ng of DNA from each sample for 8 hours at 37 °C using the restriction enzymes *PstI* and *MspI*. Samples were visually checked using a 2.5% agarose gel to ensure complete digestion. We then ligated custom adapters using T4 DNA ligase and an incubation period of three hours at 22 °C. The adapters contained a set of 96 unique 8-10 bp barcodes. Barcode sequences were generated using http://www.deenabio.com/services/gbs-adapters with at least two mutational differences between any pair of barcodes to avoid misassignment during demultiplexing. Samples were then pooled and cleaned using a QIAquick PCR purification kit. We performed 12 rounds of PCR, each cycle with 30 seconds at 98 °C, 30 seconds at 62 °C, and 30 seconds at 72 °C. Following the 12 cycles, we held for 10 minutes at 17 °C. After PCR, sets of 96 samples with unique barcodes were pooled to perform a 300-500 bp size selection using a Blue Pippen 2% agarose gel cassette. Size selection was confirmed using an ABI Bioanalyzer High Sensitivity DNA assay. Size selection and fragment analysis was performed at the Yale DNA Analysis Facility on Science Hill. We sequenced each 96-sample genomic library using one lane of Illumina HiSeq 4000 with 100 base pair single-end reads at the University of Oregon Genomics and Cell Characterization Core Facility. We inspected raw read quality using FastQC[3].

#### ddRAD Assembly

We used the *Etheostoma perlongum* genome to assemble the ddRAD dataset with iPyrad v.0.9.68[4]. After demultiplexing with iPyrad step 1, we trimmed raw reads using CutAdapt v.2.0[5]. Reads were trimmed to 100 bp and three rounds of filtering were used to remove adapter contamination and restriction cut sites ("-b TGCAG", "-a TGCAG", and "-a CCGAGATCGGAAGAGC"). Trimmed reads were then used as input for iPyrad steps 2-7 (see Table S2 for assembly parameters).

#### Genome Sequencing and Assembly

***Etheostoma perlongum* Genomic DNA Isolation --** We homogenized ~200 mg of the frozen muscle tissue by hand to a fine powder with a mortar and pestle while using liquid nitrogen to ensure the tissue remained frozen. The powder was added to a solution with 19 ml buffer G2, 10 ml buffer QBT, 30 ml buffer QC, 15 ml buffer QF, 10.5 ml isopropanol, 4 ml 70% ethanol, 38 µl RNase A (100 mg /ml), and 1 ml QIAGEN proteinase K. This solution was then incubated for 12 hours at 50 °C in a shaker set to gently rotate until all tissue powder had dissolved.

A QIAGEN Genomic-resin tip column was equilibrated with 10 ml of buffer QBT prior to adding the sample. The sample was then added to the column for gravity flow-through. We performed two washes of the column, each with 15 ml of buffer QC. Genomic DNA was eluted from the column using 15 ml of buffer QF, then precipitated by adding 10.5 ml of room-temperature isopropanol. DNA was pelleted by centrifuging for 30 minutes at 7,000 g and 4 °C. Immediately following centrifugation, the tube was rotated upside down to preserve the pellet and the buffer solution was discarded. The DNA pellet was washed with 5 ml of 70% ethanol at -20 °C, then the centrifuge step was repeated. We added 500 µl of buffer TE (pH 8.0) and incubated for 8 hours at 37 °C at a gentle rotation in a shaker to resuspend the pellet. We quantified the genomic DNA using a Quibit 3.0 and a Nanodrop. DNA quality was checked visually using a 2.5% agarose gel.

***Etheostoma perlongum* Sequencing –** For all Nanopore library preparations with the Oxford Nanopore LSK-109 ligation sequencing kit, we used wide mouth pipette tips and only mixed by inversion to avoid unnecessary shearing of large genomic DNA fragments. We also performing the optional end-repair library preparation included in the LSK-109 kit. We constructed three libraries using genomic DNA from gill tissue and two libraries using genomic DNA from muscle tissue. Libraries were sequenced using a total of four R9.4.1 flow cells. All sequencing was performed using an Oxford Nanopore Technologies MinION or GridION platform. Bases were called in real-time using Guppy v.3.0.7 (https://github.com/nanoporetech/rerio).

The 10x Genomics Chromium library was prepared by the Yale Center for Genome Analysis using the same muscle genomic DNA used for Nanopore sequencing. Pulsed-field gel electrophoresis was first used to inspect the integrity of the genomic DNA. Libraries were prepared following the standard protocol for the 10x Genomics Chromium Genome Library Kit & Gel Bead Kit v2. The library was sequenced using an Illumina NovaSeq 6000 at the Yale Center for Genome Analysis.

***Etheostoma perlongum* Genome Assembly and Scaffolding –** Prior to assembly, all Nanopore and Illumina reads were subjected to taxonomic filtering for contamination using Kraken2[6]. We compiled a custom database that included three fish genomes: *Danio rerio*[7], *Perca flavescens*[8], and *Etheostoma spectabile*[9]. All reads that were not classified as eukaryotic or that were flagged as unclassified were discarded prior to genome assembly.

Nanopore Reads were mapped against each other to construct a PAF alignment using minimap2 v.2.12 and the option "-x ava-ont"[10]. Miniasm v.0.3 was then used to assemble the reads based on the alignment from minimap2[11]. As a rough polishing step, we used minimap2 to map the Nanopore reads to the assembled contigs, then used racon v.1.4.10 to polish the assembly and generate consensus sequences[12]. We performed an additional polishing step using the unassembled 10x Chromium Illumina reads. First, we mapped the short Illumina reads to the Nanopore Miniasm assembly with bwa v.0.7.17[13], then used Pilon v.1.23 to improve base call accuracy and fix small indels and misassembles[14].

For Hi-C chromatin configuration capture sequencing, sample quality control, library preparation, and sequencing were carried out by Phase Genomics, Inc (https://phasegenomics.com/). Hi-C reads were aligned to the Nanopore contigs using the Juicer v.1.6 pipeline with default parameter values[15]. Using the output from Juicer, we ran the 3D-DNA pipeline v. 180922[16] to correct misassembles, anchor, order, and orient the contigs into scaffolds. We specified a coarse stringency threshold of 20 for each step; all other parameters were left at default values. Lastly, we used Juicebox Assembly Tools v.1.11.08 (https://github.com/aidenlab/Juicebox) to manually examine the results of 3D-DNA and curate a final scaffolded assembly for downstream analyses.

The web tool gVolante v.1.2.1 was used to assess genome assembly quality and completeness at different stages of the process by querying a BUSCO v5 ortholog dataset specific to Actinopterygii[17].

***Etheostoma nigrum* Genomic DNA Isolation, Sequencing, and Assembly** – To investigate possible structural genomic variation, we also generated a de novo genome assembly for *Etheostoma nigrum*, a closely related outgroup for *E. perlongum* and *E. maculaticeps*[18]. We extracted genomic DNA from freshly frozen muscle tissue of a single *Etheosotma nigrum* male (DJM 23, YPM ICH. 35349) using the same protocol described for *E. perlongum*. However, tissue homogenization for *E. nigrum* was achieved using a 2 mL Dounce homogenizer instead of a mortar and pestle. We prepared three Oxford Nanopore LSK-112 libraries and sequenced them using three R10.4 flow cells. Bases were called in real time using the super-accurate basecalling model in Guppy v.6.0.7 (https://github.com/nanoporetech/rerio). Prior to assembly, we used NanoFilt[19] to remove reads with quality scores $< 7$ and reads $< 500$ base pairs.

Assembly was performed using Flye v.2.9-b1774[20], with the recommended options for Q20 Nanopore reads ("--nano-hq" and "--read-error 0.03") and one iteration of polishing. We allowed Flye to select the optimal minimum overlap setting of 3,000 bp based on the Nanopore read length distribution. Flye contigs were filtered for possible contamination using the custom Kraken2 database described above. To limit the impact of haplotype heterogeneity on the assembly, we ran purge_haplotigs v.1.1.2[21] with read depth cutoffs of l=6, m=27, and h=80. We scaffolded the final set of contigs using HiC data generated by Phase Genomics, Inc and the pipelines described above. Genome assembly quality was assessed using the web tool gVolante.

### Transcriptome Sequencing and Assembly

We performed transcriptome sequencing using eight different tissues (gill, fin, gut, heart, liver, brain, eye, and testes) from the same individual used for genome sequencing (YFTC 37806, YPM ICH 033096). Tissues were immediately frozen with liquid nitrogen and stored at -80 °C. We homogenized all tissues using an MP FastPrep-24, then isolated RNA using an Invitrogen PureLink RNA Mini Kit, including the optional DNase treatment. RNA was quantified using a

Qubit 3.0 and an RNA BR Assay. We combined equimolar amounts of RNA from each tissue type for transcriptome sequencing. RNA QC, library preparation, and 150 bp paired-end Illumina sequencing was performed by NovoGene (https://en.novogene.com/).

We filtered raw reads using the *process_shortreads* program in the Stacks v.2.53 pipeline[22]. We removed reads with uncalled bases ("-c"), discarded reads with low quality scores ("-q"), and filtered to remove adapter contamination ("--adapter_1 AATGATACGGCGACCACCGAGA --adapter_2 CGTATGCCGTCTTCTGCTTG"). All reads were subjected to taxonomic filtering for contamination using Kraken2 with the same custom database used for the genomic reads. Reads that were not classified as eukaryotic or that were flagged as unclassified were discarded prior to transcriptome assembly. We assembled the transcriptome using default settings in Trinity v.2.4.0[23]. Transcriptome assembly metrics were summarized using the *TrinityStats.pl* script included with the pipeline. We used the web tool gVolante v.1.2.1 to assess transcriptome assembly quality at by querying a BUSCO ortholog dataset specific to Actinopterygii.

### Genome Annotation

We used the Maker v.2.3.10 pipeline to assign putative functional annotations to the *Etheostoma perlongum* genome assembly[24]. Evidence for Maker included the transcriptome generated in this study, set of 196,622 exons sequenced for a closely related species, *Etheostoma nigrum*[25], and protein sequences from eight other teleost species: *Etheostoma spectabile* (NCBI PRJNA606051), *Perca flavescens* (NCBI PRJNA638629), *Perca fluviatilis* (NCBI PRJNA549142), *Larimichthys crocea* (NCBI ASM74293v1), *Gasterosteus aculeatus* (Ensembl BROAD S1), *Danio rerio* (Ensembl GRCz11), and *Oryzias latipes* (Ensembl HdrR). We also included all proteins in the UniProt Swiss-Prot database (http://www.uniprit.org/; last accessed April 15, 2020). To identify repetitive elements, we generated an *E. perlongum* specific repeat library using RepeatModeler v.2.0.1, including the optional LTR structural discovery pipeline[26]. We also supplied Maker with the transposable element library included with the pipeline.

In the first annotation round, Maker soft masked repetitive elements, then used all available evidence to generate initial gene models. We used the gene models created in the first round to train the gene prediction software SNAP v.2006-07-28[27] and Augustus v.3.2.3[28]. To train SNAP, we only included gene models with a minimum length of 50 amino acids and a minimum annotation edit distance (AED) of 0.25. SNAP and Augustus were then used for a second round of *ab intio* gene prediction, again providing all protein, RNA, exon, and repeat alignments as evidence. We then performed two additional rounds of *ab intio* gene prediction with Maker, each time retraining SNAP and Augustus with gene models from the prior round. We assessed the annotation quality using the web tool gVolante v.1.2.1 by querying a BUSCO ortholog dataset specific to Actinopterygii and by calculating annotation edit distance (AED)[29]. BlastP (NCBI) was used to identify potential matches between the UniProt Swiss-Prot database (E-value threshold = 1e-6) and the *E. perlongum* predicted proteins. Lastly, InterProScan v.5[30] was used to assign putative protein domain information to the final *E. perlongum* annotations.

### Comparative Genomics

Scaffolds for *Etheostoma perlongum* were aligned to the *E. spectabile* (NCBI GCA_008692095.1), *Perca flavescens* (NCBI GCF_004354835.1), *Gadus morhua* (NCBI GCF_902167405.1), and *Oncorhynchus mykiss* (NCBI GCF_013265735.2) genomes using the CoGe SynMap2 tool (https://genomevolution.org/coge/). Coding sequences from the two genomes were aligned using the lastZ algorithm (--hspthresh 3000). DAGchainer was run using the relative gene order option; maximum distance between two genes (-D) = 20 genes; and minimum number of aligned pairs (-A) = 5 genes. Syntenic blocks were merged using Quota Align (-Dm = 0 genes). Circos v.0.69-8 was used to visualize synteny maps[31]. *Etheostoma perlongum* scaffolds were ordered according to syntenic matches with *Perca flavescens* chromosomes.

In addition, we compared the *E. perlongum* genome assembly to our de novo assembly of a close relative, *E. nigrum*. We constructed synteny maps comparing *E. nigrum* to *E. perlongum* using the methods described above. We also used nucmer v.4.0.0[32] to perform genome-genome alignment with default options. Using a custom R script, we visualized the alignment dotplot, discarding alignments < 5000 bp. Lastly, we examined an *E. nigrum* contig that spanned the *E. perlongum* chromosome 9 inversion breakpoint. Nucmer was used to align the *E. nigrum* contig to *E. perlongum* chromosome 9, and visualization was performed with the R package RIdeogram v.0.2.2[33].

**Identification of Linkage Disequilibrium Clusters**

For LDna[34], the choice of two parameters, φ and |E|min, influences which clusters of high LD markers are considered outliers. |E|min specifies the minimum number of edges for a cluster, while φ specifies a minimum LD threshold for identifying outliers. Based on recommendations from the LDna authors (https://github.com/petrikemppainen/LDna/), we explored several different |E|min thresholds, eventually selecting |E|min = 20 to avoid noise from smaller networks of tight physical linkage. Following[35], we ran several analyses for each chromosome, starting with φ = 0 and incrementing the value of φ by 1 until no LD outlier clusters were detected. When clusters at different φ shared SNPs, we retained the cluster with the fewest number of SNPs and highest intra-cluster LD. As we sought to identify large outlier clusters with high LD that could represent chromosomal inversions, we retained SOCs containing more than 30 SNPs and with median intra-cluster LD greater than 0.3.

**Population Structure Analyses**

While filtering removed SNPs with large amounts of missing data, some individuals in our dataset still contained large proportions of missing data, presumably due to technical errors during library preparation. Therefore, we used the *impute* function in the R package LEA v.2.6.0 to replacing missing genotypes with a random value weighted by the observed genotype probabilities[36].

As a first pass at examining population structure, PCA was performed on the ddRAD SNPs using the R package stats v.4.0.0 (R Core Team 2020). We then applied a model-based clustering approach used sparse non-negative matrix factorization (sNMF) to estimate individual ancestry coefficients with the R package LEA. We incremented the number of genetic clusters (K) from one to ten, performing 10 repetitions for each K value. We set the regularization parameter (alpha) to 10, the tolerance to 0.00001, and the percentage of masked genotypes for cross-entropy validation to 0.05. For each K value, the replicate with the minimum cross-entropy score was selected for further visualization and comparison. A cross-validation approach using cross-entropy criteria was performed to determine the optimal number of genetic clusters to describe the data.

Due to limitations in the *snapclust* method[37], we reduced the size of our SNP dataset by enforcing a stricter missing data threshold with VCFTools ("--max-missing 0.96"). We first used the *find.clusters* function to implement K-means clustering, specifying K=2 and using 10 million search iterations for each of 1,000 random starting centroids. We used the K-means clustering results as input for the s*napclust* function, performing 1,000 runs of the EM maximum likelihood algorithm with an upper limit of 10 million search iterations. We performed two sets of *snapclust* analyses: allowing only F1 hybrids ("hybrid.coef=0.5") and allowing F1 hybrids plus backcrosses ("hybrid.coef=c(0.25, 0.5)").

**Demographic Modeling**

To construct a SFS containing monomorphic sites, we converted the iPyrad "loci" file to VCF format using a custom python script. We filtered this dataset using VCFTools[38] to remove sites with >10% missing data ("--max-missing 0.9") and sites with more than two alleles ("--max-alleles 2"). To account for missing data, a down-projected SFS[39] was constructed from the filtered VCF file using easySFS (https://github.com/isaacovercast/easySFS), retaining 50, 42, and 76 haploid samples for *E. perlongum* in Lake Waccamaw, the hybrid population, and *E. maculaticeps* in the Waccamaw River, respectively.

Estimation of demographic parameters for the three models required either a fixed mutation rate or a fixed root age. Since there are no genome-wide mutation rate estimates for darters and the root age was a parameter of interest, we ran a series of simulations with fastsimcoal2 v.2.6.0.3 [40], fixing the root age at 1,000 generation intervals between 1,000 and 32,000 generations. The upper limit was based on sediment core estimates of the maximum age of Lake Waccamaw[41]. For each model at each fixed root age, we performed 40 sets of simulations with fastsimcoal2, running 100,000 simulations for 40 cycles of the ECM likelihood optimization algorithm. We drew initial values from a log-uniform distributions for the $N_e$ parameters (min=100, max=100,000), migration rate parameters (min=1e-8, max=0.5), and mutation rate parameter (min=1e-12, max=1e-3). Note that the minimum and maximum values were not hard boundaries for these parameters. We drew initial values for the hybrid population origin time from a uniform distribution (min=1, max=root age – 1), imposing a hard upper boundary. We drew initial values for the starting admixture proportion of the hybrid population from a uniform distribution (min=0, max=1), again imposing a hard upper boundary. We compared model fit by calculating AIC scores.

For the best-fit demographic model, we generated 100 non-parametric bootstrap replicates using a custom script to resample 1 Mb blocks from the original VCF file. For each bootstrap replicate, we performed analyses as outlined above, but specifying the initial parameter values using the final parameter values from the best-fit run. We then calculated the 95% highest density interval for each parameter using the R package *bayestestR* v.0.7.5[42].

### Genetic Geographic Clines

Allele frequency clines were fit using the R package HZAR v.0.2.5[43]. For all localities, we calculated river distance from the small dam separating Lake Waccamaw and the Waccamaw River (34.260890, -78.523142) using the R package riverdist v.0.15.1[44]. The cline models all have two important parameters in common: the cline center and the cline width. For the cline center, zero km is defined as the Lake Waccamaw outlet. Cline center was restricted to values between -10 and 320 km, the limits of our observed data. We also restricted the cline width, deltaL, and deltaR parameters to values between 0 and 330 km. For each SNP, following an initial run to burn-in each model, we performed three replicate MCMC optimizations per model using the function *hzar.doChain.multi*. Starting parameter values were randomized for each replicate run. Each MCMC optimization was run for one million generations with 100,000 generations of burn-in and a thinning interval of 100 generations. We then performed model selection using AICc scores.

### Genetic Outlier Detection

We first performed a genome scan for selection to detect outlier loci using the R package pcadapt v.4.3.3[45]. Briefly, this approach computes z-scores based on Mahalanobis distances from a regression of SNPs versus the first K principal components. Based on the results of scree plots for preliminary runs, we retained only a single PC axis, which differentiated the lake and river samples. We then performed the Benjamini-Hochberg procedure to control the false discovery rate with alpha = 0.0001 using the *p.adjust* function in the R core package stats v.4.0.0 (R Core Team 2020).

In addition, we performed two genome scans for selection that utilize the fixation index, $F_{ST}$[46]. First, we used the *populations* program in the Stacks v.2.53 pipeline[22] to perform AMOVA $F_{ST}$ estimation for each SNP. We supplied the program with the same VCF file used with pcadapt. We identified outliers as SNPs above the 99$^{th}$ percentile of differentiation. In addition, to examine whether the $F_{ST}$ results were driven by sex-ratio sampling biases, we also assembled male and female SNP subsets. We confidently identified 5 females and 7 males from the Waccamaw River and 6 females and 5 males from Lake Waccamaw. Sample sizes were low because many of the specimens were too small to accurately sex. We estimated $F_{ST}$ for the male and female subsets with the Stacks pipeline[22].

To complement the AMOVA $F_{ST}$ estimates, we also performed Bayesian estimation of $F_{ST}$ using BayeScan v.2.1[47]. We first converted the VCF file to BayeScan input format using PDGSpider v.2.1.1.5[48]. We specified a prior odds ratio of 10 and ran four independent analyses, each with 20 pilot runs of 5,000 generations, 50,000 generations of burnin, then 50,000 sampling generations with a thinning interval of 10. We assessed convergence and mixing of the MCMC replicate runs, ensuring ESS values were greater than 200 using the R package CODA v.0.19-3[49]. We combined posterior estimates from all four runs. SNPs were considered outliers if their FDR q-value was less than 0.05.

We also estimated $F_{ST}$ for 1 Mb windows with a slide size of 100 Kb. We used a large window size because ddRAD SNPs are sparsely distributed across the genome. To calculate diversity statistics for these windows, we used a set of python scripts from Simon Martin (https://github.com/simonhmartin/genomics_general). To estimate absolute divergence ($D_{XY}$)[50,51] and nucleotide diversity ($\pi$)[50], we required an input VCF file containing variant and invariant sites. Therefore, we converted the iPyrad "loci" format to VCF using a custom python script. We then removed sites with more than two alleles ("--max-alleles 2") and removed sites with more than 10% missing data ("--max missing 0.9") using VCFTools. The filtered dataset contained 2,270,061 total sites. We next converted the VCF file to a geno file using the *parseVCF.py* script and estimated $F_{ST}$, $D_{XY}$, and $\pi$ (lake and river) for sliding windows using the *popgenWindows.py* script. We identified $F_{ST}$ and $D_{XY}$ outliers as regions above 99th percentile of differentiation.

### Gene Ontology Enrichment Analysis

We examined three genomic windows on chromosome 9 with elevated lake-river divergence. The $F_{ST}$ plateau (Fig. 4A, chromosome 9:24,641,313..33,628,684) was defined as the region containing SNPs identified as outliers by at least two out of three methods, plus 10 Kbp on either side. $D_{XY}$ peak 1 (Fig. 4A, chromosome 9:24,900,001..25,900,000) and $D_{XY}$ peak 2 (Fig. 4A, chromosome 9:29,400,001.. 31,100,000) were regions within the $F_{ST}$ plateau that were also idented as $D_{XY}$ outliers. These genomic regions contained 393, 10, and 83 genes, respectively. For BlastX searches, we retained the first 20 hits with an E-value threshold of 1e-6. We discarded hits shorter than 20 amino acids or with less than 70% sequence similarity and retained the hit for each gene with the highest HSP.

### Morphological Analyses

Photographs of the left lateral side of specimens were captured using a Nikon D5100 and a 60 mm macro lens. Specimens were pinned to a dissecting mat to ensure that all landmark positions were visible and to straighten bent specimens when possible. We supplemented our photographs with previously published images[52].

To correct for specimen warp during preservation, we used five landmarks along the lateral line with the *unbend* module in tpsUtil v.1.78[53]. We also corrected for jaw position by rotating the lower jaw landmark using the *fixed.angle* function in the R package geomorph v.3.3.1[54]. Finally, we aligned and resized all specimen landmarks using a generalized procrustes analysis as implemented by the geomorph function *gpagen*[54].

Caudal length was defined as the distance between the narrowest portion of the caudal peduncle and the origin of the caudal fin. We used the *interlmkdist* function in the R package geomorph to extract these measurements from specimens prior to using tpsUtil *unbend* because this module rescales landmarks. The four linear measurements were corrected for body size by taking the residuals of a linear regression against standard length (measured from tip of the snout to the origin of the caudal fin).

To achieve reasonable samples sizes at each locality, we used a subset of our data consisting of 185 individuals from nine localities spanning the lake and river. We used the HZAR function *hzar.doMorphoSets* to transform the morphological data using Bernoulli trials to values between 0 and 1 that were appropriate for geographic cline model fitting. Since the morphological sampling localities were more geographically restricted, we constrained the cline center to values between -10 and 100 km and constrained the cline width, deltaL, and deltaR parameters to values between 0 and 110 km.

In addition to the geometric landmark morphometrics analysis, we collected data from 86 individuals (23 Lake Waccamaw, 25 hybrid zone, and 38 Waccamaw River) for 16 meristic traits used in traditional ichthyological taxonomy: lateral line scales, pored lateral line scales, transverse scale rows, scales around the caudal peduncle, anal fin spines, anal fin rays, scales above the lateral line, scales below the lateral line, first dorsal fin spines, second dorsal fin rays, pectoral fin rays, percent cheek squamation, percent opercle squamation, percent nape squamation, percent breast squamation, and percent belly squamation. The numbers of scale rows and fin elements were determined from each specimen as outlined in[55,56], with the exception of the number of transverse scale rows, which was counted as described by[57]. Delineation of areas for squamation follow[58]. We also quantified two pigmentation patterns: the number of lateral bands and dorsal saddles. Counts are provided in Dataset S3.

Finally, we also collected linear measurements of morphology from micro computed tomography (µCT) scans. We scanned a subset of specimens that included 18 individuals: eight from the lake, six from the river, and four from the hybrid zone. Scans were done on a Skyscan 1173 (Bruker microCT, Kontich, Belgium) at the Museum of Comparative Zoology at Harvard University using resolutions of 14-22 µm per pixel and a range of scan parameters (40-65 kV, 123-200 µA, 500-1000 ms exposure). Scans were reconstructed using NRecon (Bruker microCT) and the resulting slices were loaded into 3DSlicer v4.11.0-2020-07-18[59] for visualization, segmentation, and data collection.

A variety of linear measurements were done on the skeleton of each scanned individual, largely following previous work on comparative darter morphology[60]. These included 10 linear measurements and two ratios which combined two linear measurements (Fig. S6):

1) The standard length; measured from the anterior-most tip of the snout and jaws to the posterior end of the hypural plate at the base of the caudal fin.
2) The head length; measured from the joint between the neurocranium and the first vertebral centra and the anterior-most tip of the snout.
3) The hyoid length; measured on a single side from the anterior-most tip of the hyoid to the posterior-most tip of the hyoid, including the hypohyal, anterior ceratohyal, and posterior ceratohyal bones, but not the small interhyal.
4) The length of the dentigerous premaxilla; measured as the anterior-most part of the toothed arm of the premaxilla to the posterior-most point.
5) The buccal cavity length; as measured from the anterior-most point of the dentary symphysis to the anterior-most point of the hyoid symphysis.
6) The gape width; measured between the two dorsal-most points of the coronoid processes of each side of the dentary.

7) The lower jaw out lever; measured from the anterior tip of the dentary to the center of the quadrate-articular joint.
8) The lower jaw closing in-lever; measured from the insertion of the adductor mandibulae on the posterior slope of the coronoid process of the articular to the center of the quadrate-articular joint.
9) The lower jaw closing in-lever at the tip of the coronoid process; measured as in #8 but from the dorsal tip of the coronoid process to the center of the quadrate-articular joint.
10) The lower jaw opening in-lever: measured from center of the insertion of the interoperculomandibular ligament on the retroarticular process of the articular to the center of the quadrate-articular joint.
11) The lower jaw opening ratio; calculated by dividing the lower jaw opening in-lever (#10) by the lower jaw out lever (#7)
12) The lower jaw closing ratio; calculated by dividing the closing in-lever (#8 or #9) by the out lever (#7). Note that because we made two different but highly correlated measurements of the closing in-lever (#8 and #9), we can calculate this ratio in two ways. Unless otherwise noted, we used #8, to be consistent with[60].

With a few exceptions, these measurements are all related to the feeding anatomy of fishes, and we would expect to see differences in some of these measurements with changes in a species ecology and diet. The two lever ratios – closing and opening jaw ratios – are also biomechanical measurements of lower jaw performance along a force-mobility axis[61]. Low values of closing or opening lever ratio are characterized as "mobility" modified, indicating enhanced movement at the out-lever given a similar movement of the in-lever. These low values are associated with higher reliance on suction feeding compared to biting. The opposite is true of high values of opening or closing lever ratios, which are described as "force" modified and are associated with increased biting during feeding.

For analysis, all linear measurements (but not the two ratios) were size-corrected by dividing by standard length. Data for these linear measurements is given in Dataset S4.

We analyzed morphometric, osteological, and meristic trait differences between Lake Waccamaw (*E. perlongum*), the hybrid zone in the lake outlet (locality 6, Fig. 1), and the Waccamaw River (*E. maculaticeps*) using ANOVA and post-hoc Tukey HSD tests with R the stats package v.4.0.0 (R Core Team 2020). For osteological and meristic traits, we also examined correlation with genomic ancestry coefficients estimated by LEA for the subset of individuals included in the ddRAD dataset. For the osteological traits, the ddRAD subset contained six individuals from Lake Waccamaw, two individuals from the hybrid zone at the lake outlet, and six individuals from the Waccamaw River. For the meristic traits, the ddRAD subset included eight individuals from Lake Waccamaw, eight individuals from the hybrid zone at the lake outlet, and seven individuals from the Waccamaw River. We performed linear regression using the R the stats package.

**Diet Analyses**

We assessed the trophic niche of Lake Waccamaw and Waccamaw River populations by collecting diet data from gut contents of specimens of *Etheostoma perlongum* (YPM ICH 28448, 32699, and 33096; NCSM 6918, 8927, and 56309), hybrid (YPM ICH 31397; NCSM 51791), and *E. maculaticeps* (YPM ICH 28323 and 28947; NCSM 65497). The digestive tract, from between the oesophagus to the anal vent, were removed from fish, and contents of the foregut were identified to ordinal or family level, where possible using taxonomic keys[62–64]. The number of individuals of each prey taxon in each stomach were quantified based on the number of head capsules or other uniquely identifiable body part observed.

We estimated multivariate trophic ecospace using Nonmetric Multidimensional Scaling (NMDS). NMDS was performed based on Brays-Curtis dissimilarity with the R package *vegan* v. 2.5-7[65]. We performed analyses of two- or three- dimensional ordinations and based our interpretation on the ordination with the lower stress score. The first two dimensions were used for visualization of *E. perlongum* and *E. maculaticeps* trophic ecospace. Association with trophic resources was also visualized using the *plotweb* function the R package bipartite v. 2.16[66,67] to construct a bipartite network to compare the percent frequency occurrences of each major prey category in the diets of *E. perlongum* and *E. maculaticeps*. Prey identifications were collapsed to order-level for Coleoptera, Ephemeroptera, Isopoda, and Trichoptera and class-level for bivalves because some fish stomachs contained partial prey that did not allow positive identification to lower taxonomic levels.

### SI RESULTS

#### *Etheostoma nigrum* genome assembly

In order to examine putative structure genomic variation, we used Nanopore long reads scaffolded with a Hi-C contact map to produce a de novo chromosome-level genome assembly for *Etheostoma nigrum*, a close relative of *E. perlongum.* We sequenced 39.56 Gb of Nanopore reads with a median read length of 5,470 bp, a read length N50 of 10,570 bp, and a mean read quality score of 16.6, representing approximately 46X genomic coverage. The *E. nigrum* genome assembly is 764 Mb in length, comprised of 6,010 contigs (N50 = 0.8 Mb, L50=215) contained in 3,559 scaffolds (N50 = 30.2 Mb, L50 = 12). The 24 longest scaffolds account for 93.4% of the total assembly length and exhibit a 1:1 relationship with the *E. perlongum* chromosomes (Fig. S1). Although we did not annotate the *E. nigrum* genome, 98.0% of BUSCO Actinopterygii orthologs are represented as complete sequences in the assembly.

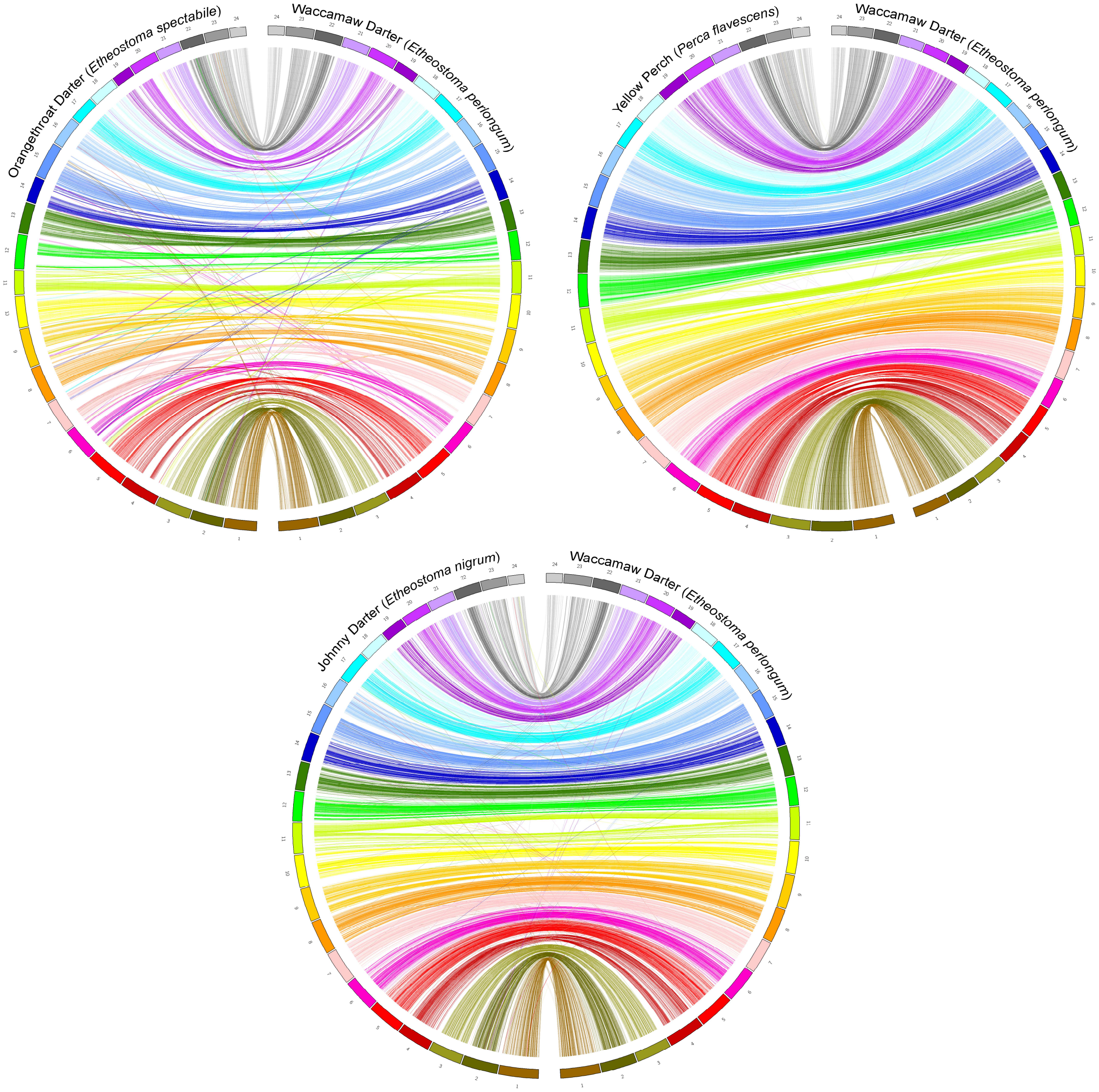

**Fig. S1.** Circos plot comparing gene synteny with the *Etheostoma perlongum* genome. Each link represents a syntenic gene pair, colored by *E. perlongum* scaffold.

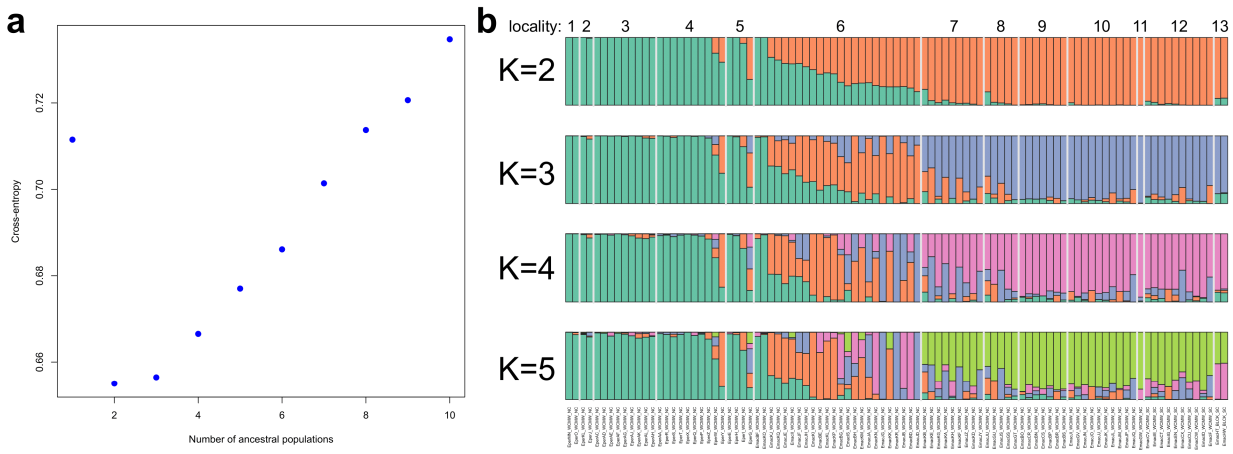

**Fig. S2.** Population genetic clustering. a) Minimum sNMF cross-entropy scores from 10 replicate runs of each K value. b) Genetic clustering analyses with 2 to 5 genetic clusters (K). Each vertical bar represents a sample, bar colors represent admixture coefficients estimated by sNMF. Sampling sites are delimited by vertical white bars. Locality numbers as in Figure 1A.

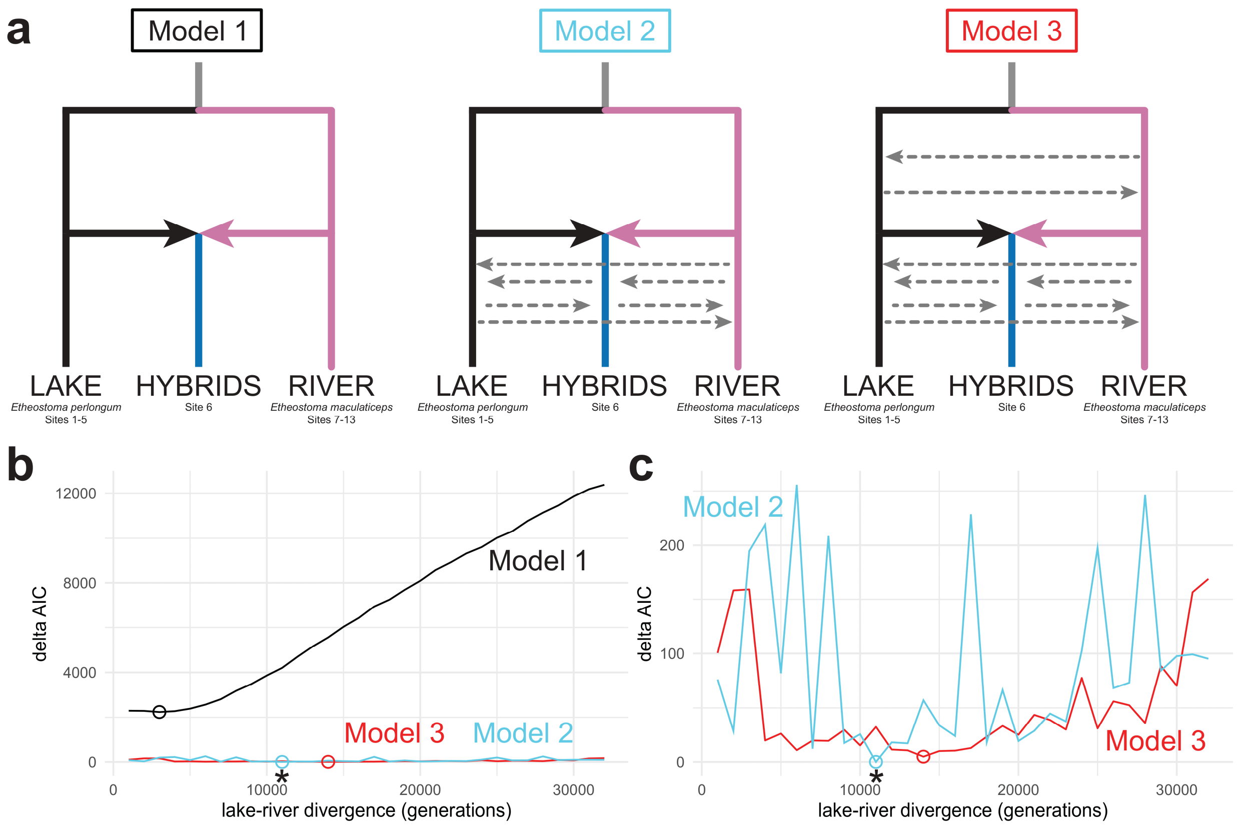

**Fig. S3.** Demographic modeling with fastsimcoal2. A) Three tested models with a hybrid population formed by admixture between river and lake with migration events visualized as dotted grey lines. Model 1: no migration. Model 2: migration only after hybrid zone is formed. Model 3: continuous migration after initial divergence. B) ΔAIC values for each model with different fixed lake-river divergence times. Divergence times range from 1,000 to 32,000 in increments of 1,000 generations. Run with the best AIC score for each model is circled. C) Same as in B, but only displaying models 2 and 3.

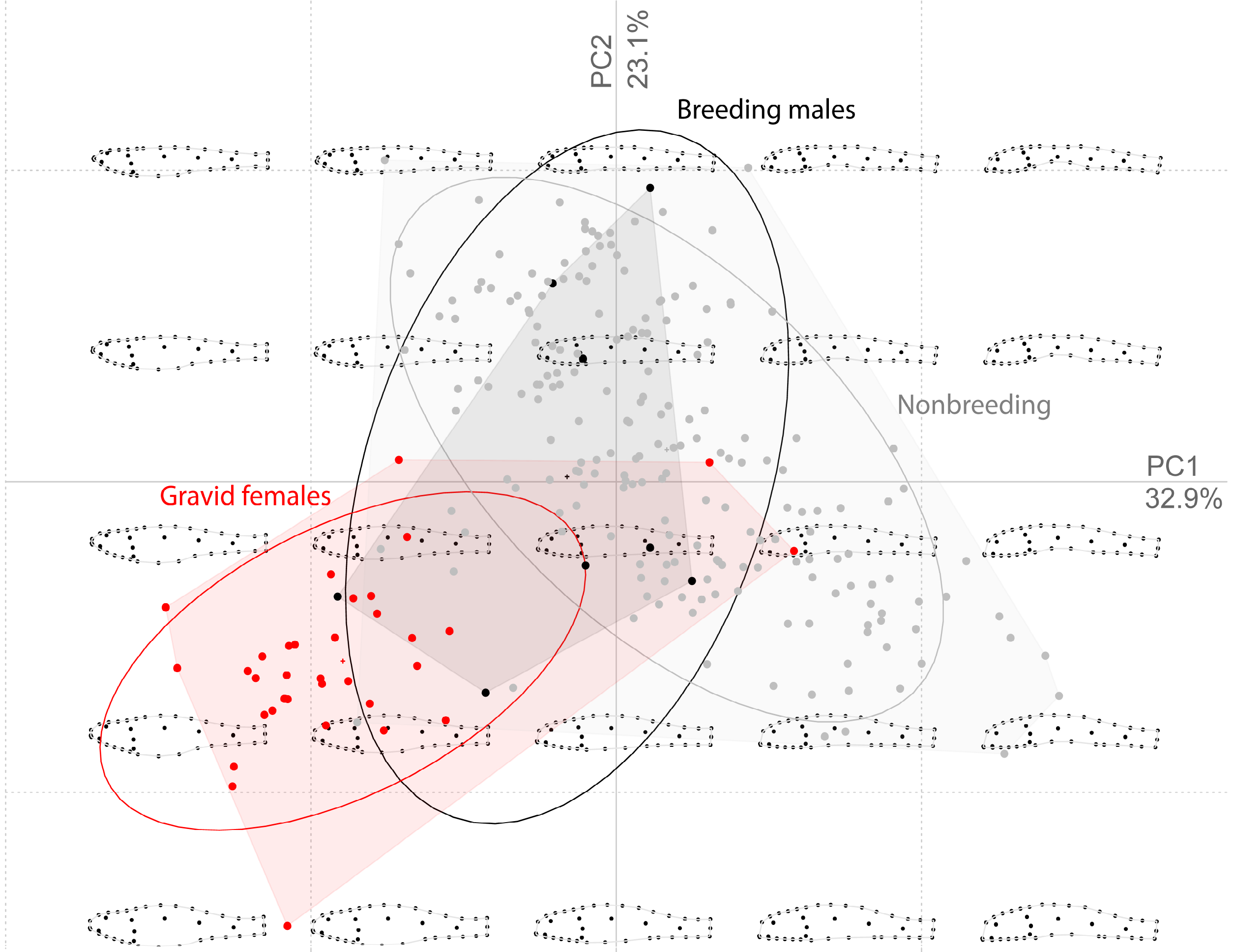

**Fig. S4.** 2D-body shape morphospace showing first and second principal components of the geometric morphometric analysis. Filled regions are convex hulls with 95% credible interval ellipsoids around the mean for each group. Gray background outlines represent body shapes in different regions of morphospace.

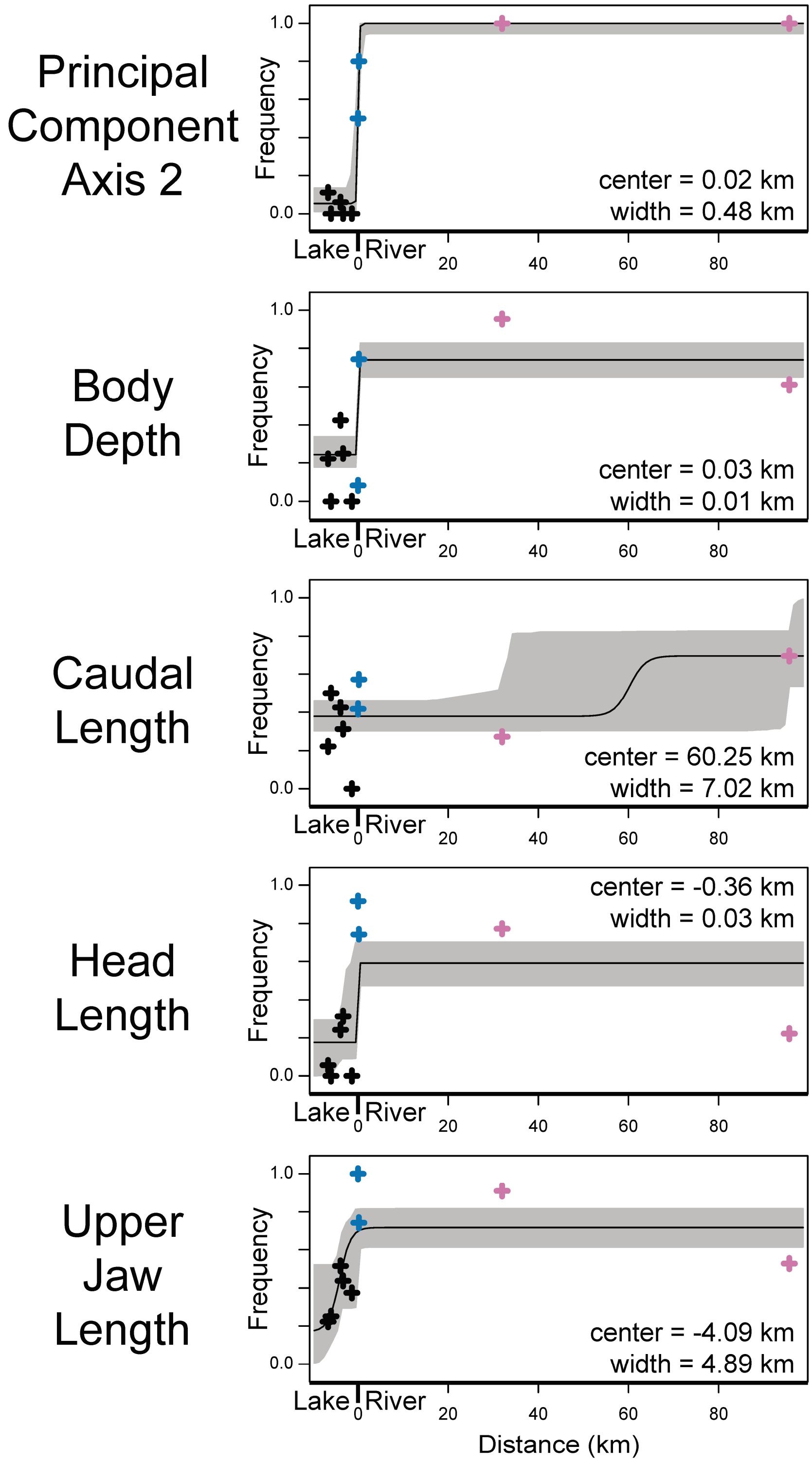

**Fig. S5.** Geographic clines in morphological traits. All traits (except PC2) have been transformed using Bernoulli trials to values between 0 and 1. Black lines indicates the maximum likelihood cline, gray areas represent 95% credible cline region. Pink points indicate *Etheostoma maculaticeps*, blue points indicate hybrids, and black points indicate *E. perlongum.*

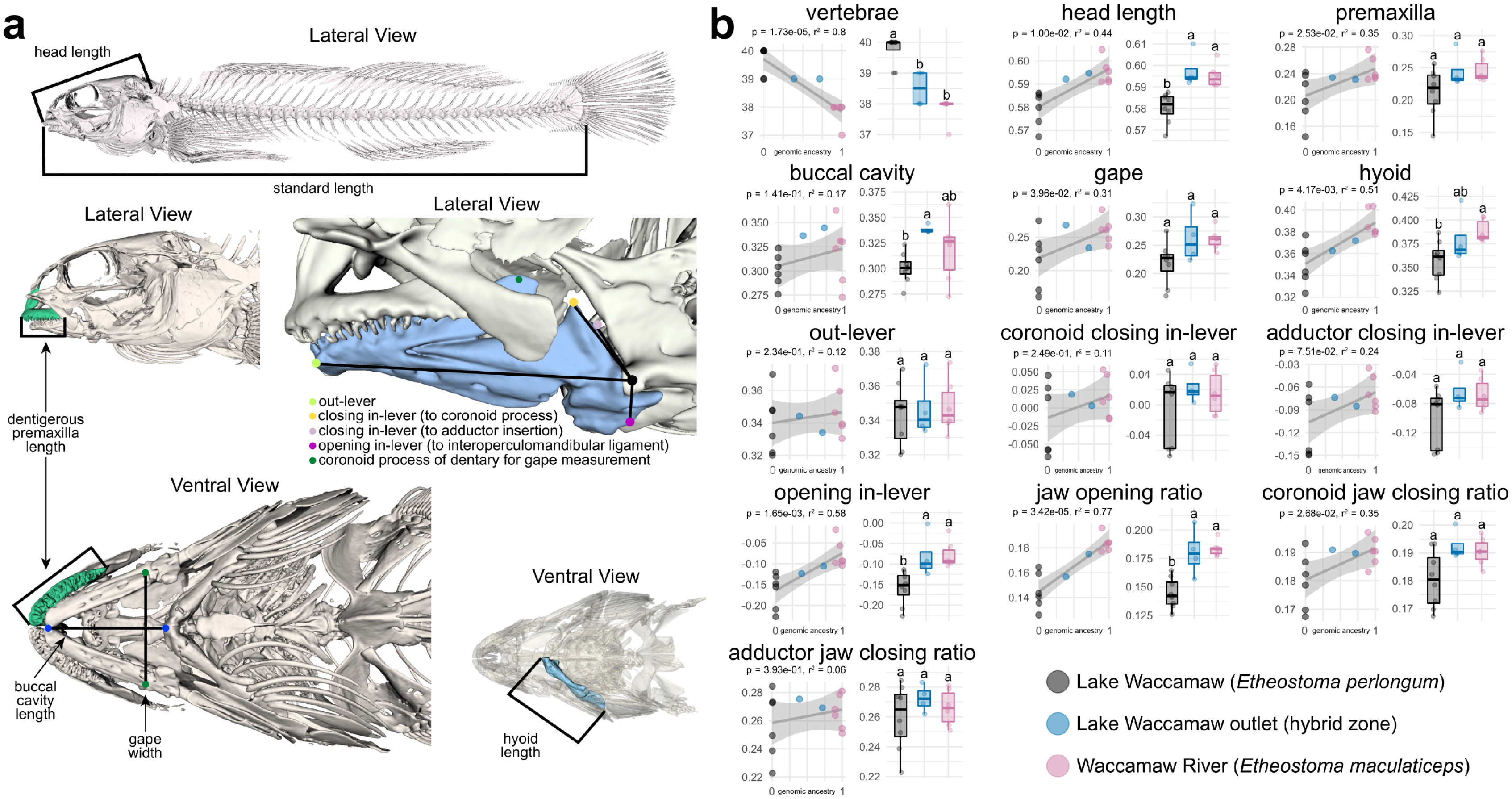

a
Lateral View
head length
standard length
Lateral View
Lateral View
dentigerous premaxilla length
out-lever
closing in-lever (to coronoid process)
closing in-lever (to adductor insertion)
opening in-lever (to interoperculomandibular ligament)
coronoid process of dentary for gape measurement
Ventral View
buccal cavity length
gape width
Ventral View
hyoid length
b
vertebrae
p = 1.73e-05, r² = 0.8
head length
p = 1.00e-02, r² = 0.44
premaxilla
p = 2.53e-02, r² = 0.35
buccal cavity
p = 1.41e-01, r² = 0.17
gape
p = 3.96e-02, r² = 0.31
hyoid
p = 4.17e-03, r² = 0.51
out-lever
p = 2.34e-01, r² = 0.12
coronoid closing in-lever
p = 2.49e-01, r² = 0.11
adductor closing in-lever
p = 7.51e-02, r² = 0.24
opening in-lever
p = 1.65e-03, r² = 0.58
jaw opening ratio
p = 3.42e-05, r² = 0.77
coronoid jaw closing ratio
p = 2.68e-02, r² = 0.35
adductor jaw closing ratio
p = 3.93e-01, r² = 0.06
genomic ancestry
Lake Waccamaw (Etheostoma perlongum)
Lake Waccamaw outlet (hybrid zone)
Waccamaw River (Etheostoma maculaticeps)

**Fig. S6.** Analyses of osteological data. A) Linear measurements of morphology from micro computed tomography (μCT) scans. B) Linear regressions of osteological traits versus genomic ancestry coefficients estimated by SNMF (Fig. 1D). Points are colored by sampling location (lake, outlet, or river). Boxplots of for each trait grouped by locality (lake, outlet, or river), with compact letter displays of Tukey HSD tests indicated above each plot. Boxplots center line shows the median; box limits show 25th and 75th quartiles; whiskers show 1.5x interquartile range; all data points shown.

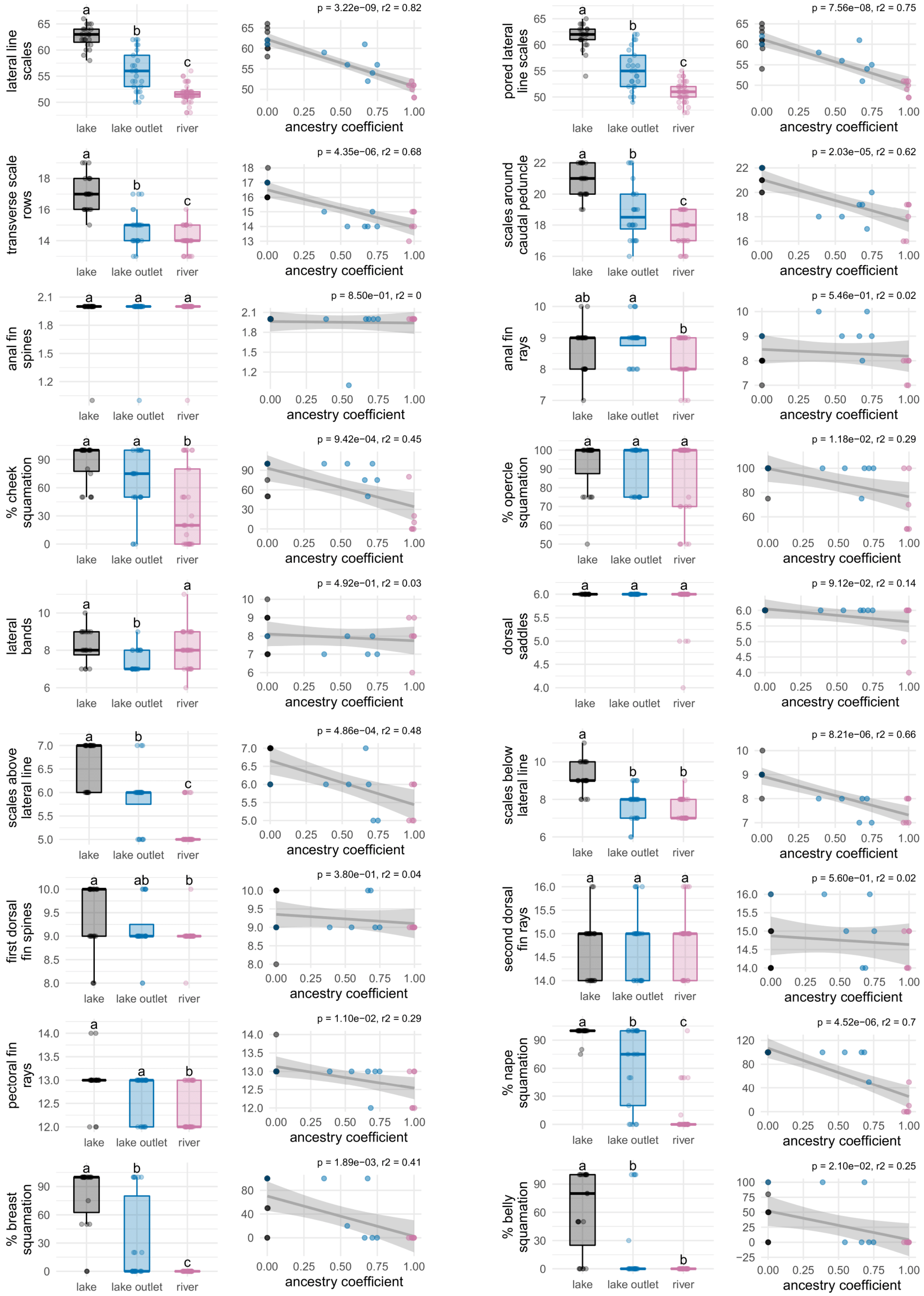
lateral line scales
p = 3.22e-09, r2 = 0.82
pored lateral line scales
p = 7.56e-08, r2 = 0.75
transverse scale rows
p = 4.35e-06, r2 = 0.68
scales around caudal peduncle
p = 2.03e-05, r2 = 0.62
anal fin spines
p = 8.50e-01, r2 = 0
anal fin rays
p = 5.46e-01, r2 = 0.02
% cheek squamation
p = 9.42e-04, r2 = 0.45
% opercle squamation
p = 1.18e-02, r2 = 0.29
lateral bands
p = 4.92e-01, r2 = 0.03
dorsal saddles
p = 9.12e-02, r2 = 0.14
scales above lateral line
p = 4.86e-04, r2 = 0.48
scales below lateral line
p = 8.21e-06, r2 = 0.66
first dorsal fin spines
p = 3.80e-01, r2 = 0.04
second dorsal fin rays
p = 5.60e-01, r2 = 0.02
pectoral fin rays
p = 1.10e-02, r2 = 0.29
% nape squamation
p = 4.52e-06, r2 = 0.7
% breast squamation
p = 1.89e-03, r2 = 0.41
% belly squamation
p = 2.10e-02, r2 = 0.25
lake
lake outlet
river
ancestry coefficient

**Fig. S7.** Analyses of meristic data. Compact letter displays of Tukey HSD tests indicated above each box plot. Boxplots center line shows the median; box limits show 25th and 75th quartiles; whiskers show 1.5x interquartile range; all data points shown. Linear regressions genomic ancestry coefficients estimated by SNMF (Fig. 1D) versus meristic trait values. Points are colored by sampling location (lake, outlet, or river).

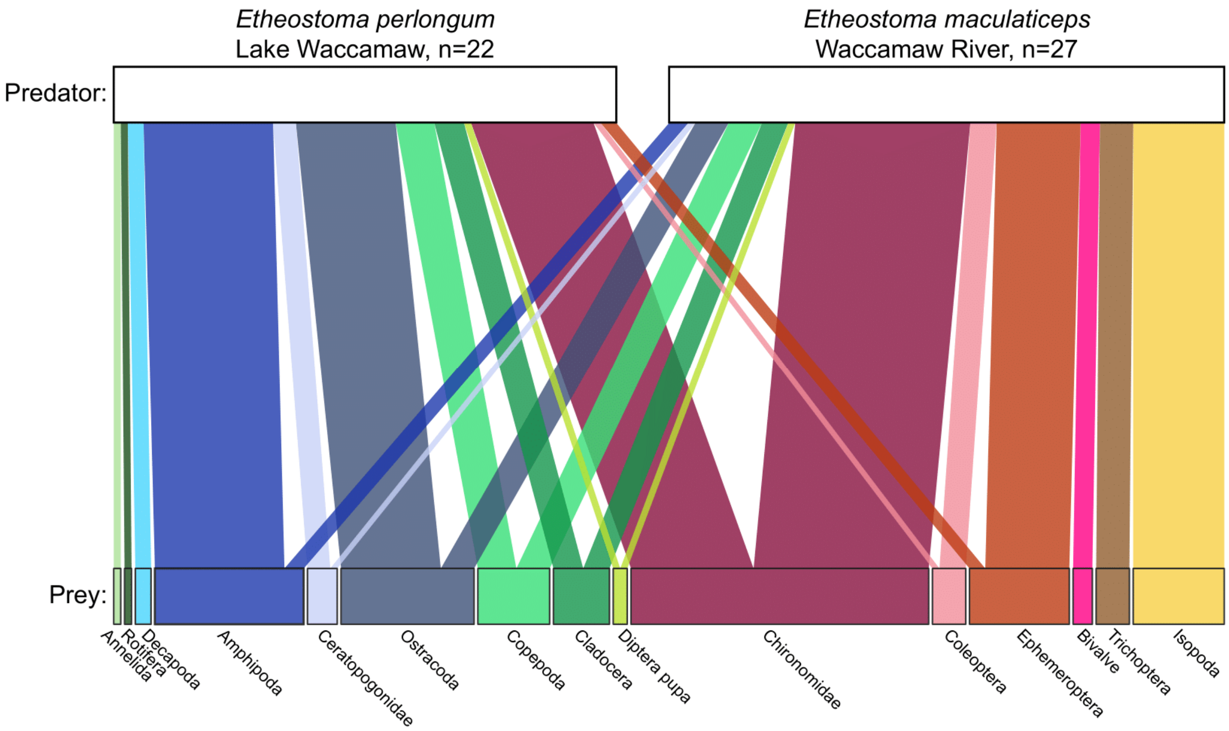

**Fig. S8.** Bipartite network of *Etheostoma perlongum* and *E. maculaticeps* diets. Width of network edges is proportional to the percent frequency occurrence of each prey category.

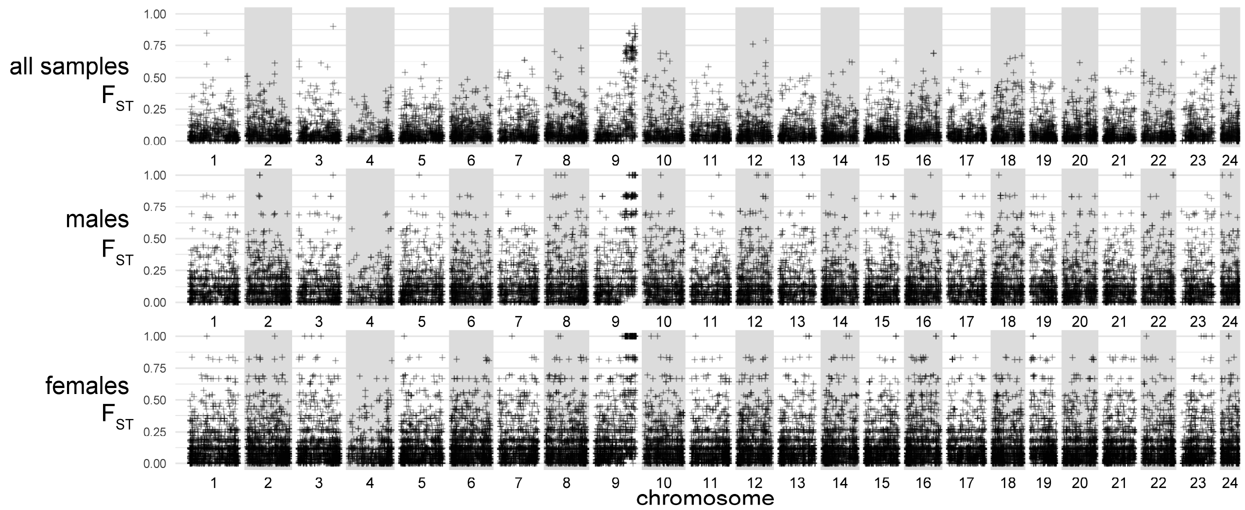

**Fig. S9.** SNP $F_{ST}$ estimates. All individuals versus males (n=12) and females (n=11) separately.

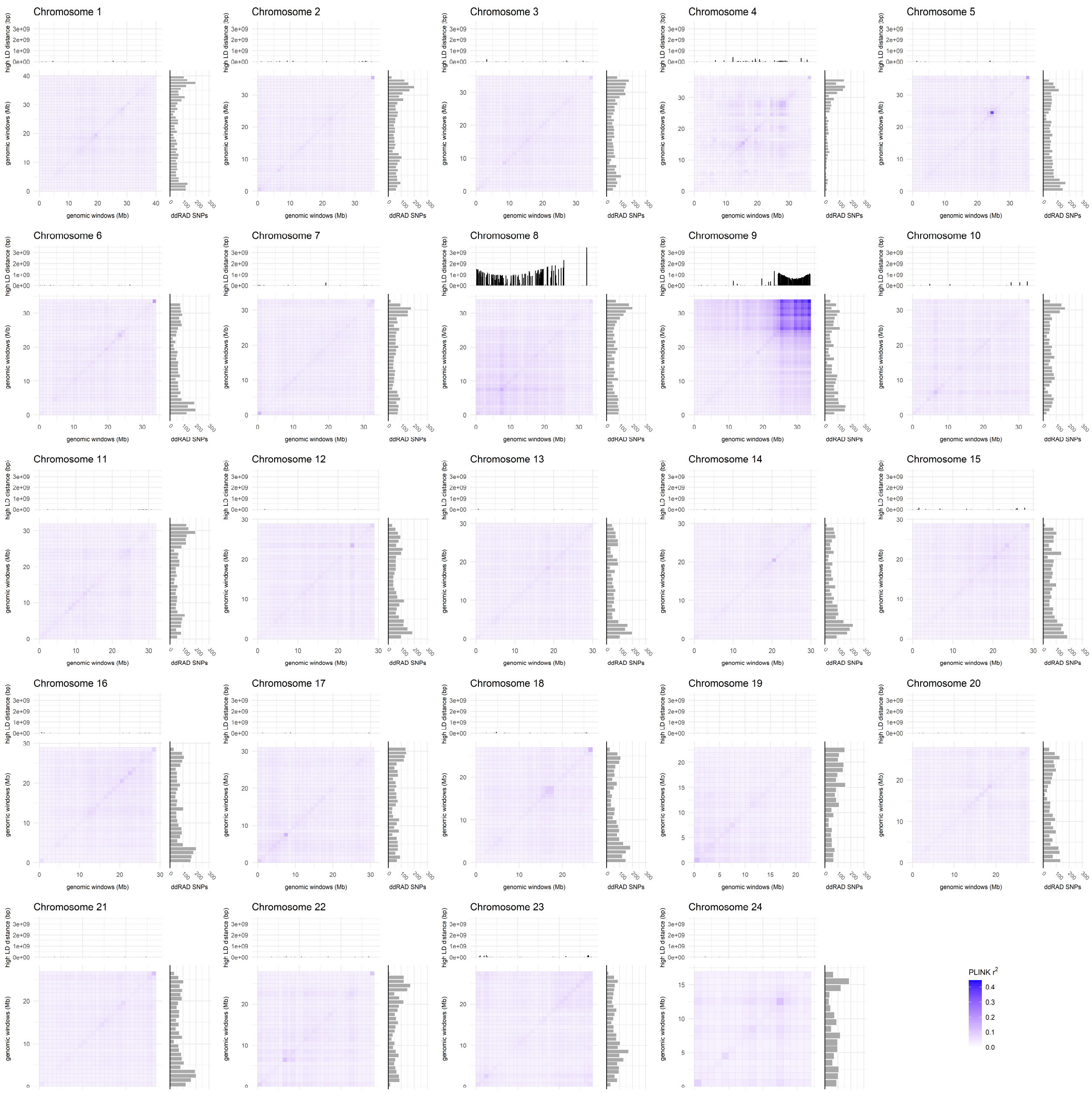

**Fig. S10.** Linkage disequilibrium (LD) heatmap in 1 Mbp windows for all lake and river samples. Bar plots on the top margins show per-SNP linkage metric calculated as the sum of distances between SNPs in high LD (R2 > 0.8). Histograms on the right margins show the distribution of ddRAD SNPs in 1 Mbp windows along chromosome 9.

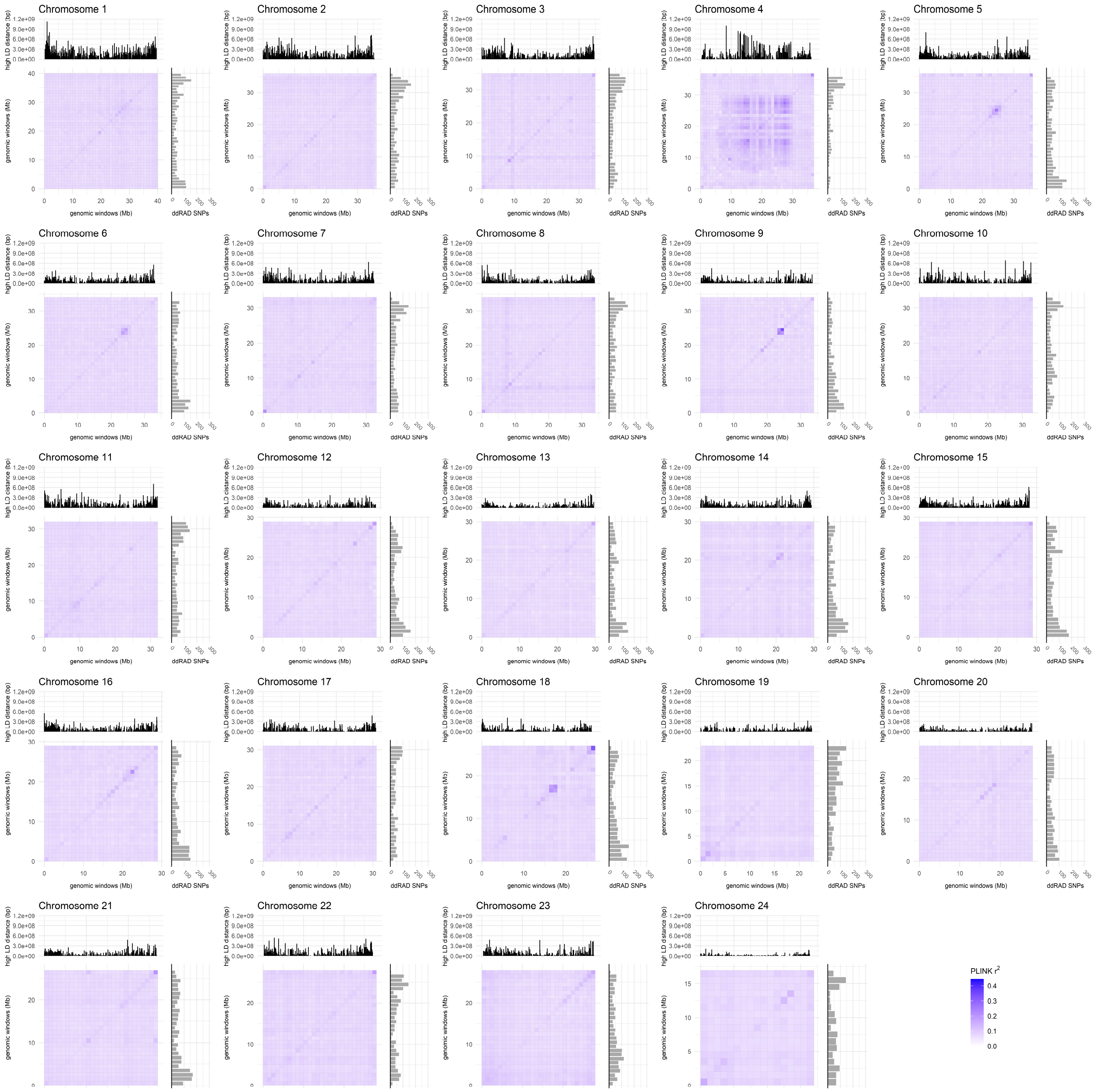

**Fig. S11.** Linkage disequilibrium (LD) heatmap in 1 Mbp windows for only samples with at least 75% of SOC-1 SNPs homozygous for the reference (lake) chromosome 9 inversion genotype. Bar plots on the top margins show per-SNP linkage metric calculated as the sum of distances between SNPs in high LD (R2 > 0.8). Histograms on the right margins show the distribution of ddRAD SNPs in 1 Mbp windows along chromosome 9.

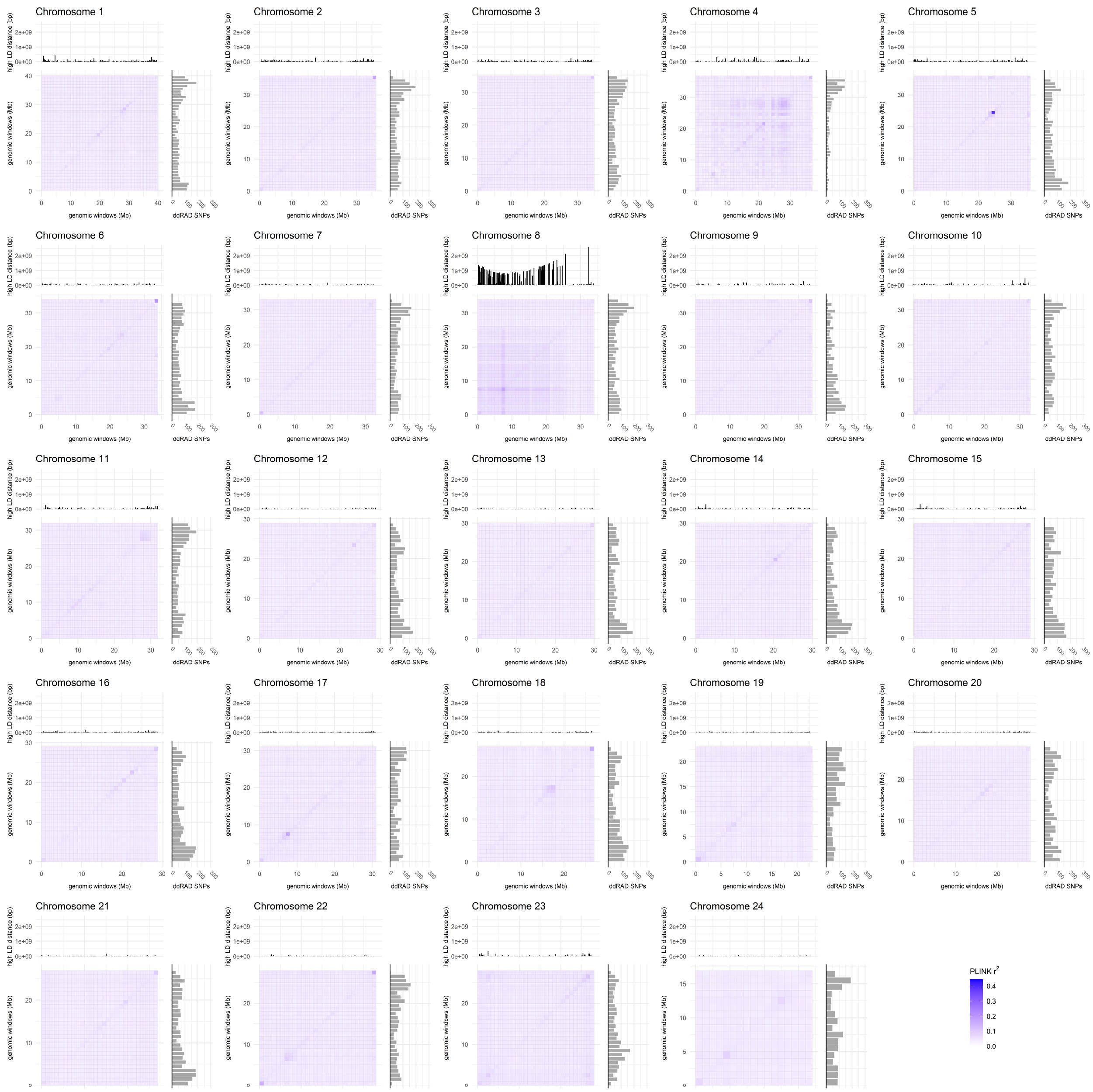

**Fig. S12.** Linkage disequilibrium (LD) heatmap in 1 Mbp windows for only samples with at least 75% of SOC-1 SNPs homozygous for the alternate (river) chromosome 9 inversion genotype. Bar plots on the top margins show per-SNP linkage metric calculated as the sum of distances between SNPs in high LD ($R2 > 0.8$). Histograms on the right margins show the distribution of ddRAD SNPs in 1 Mbp windows along chromosome 9.

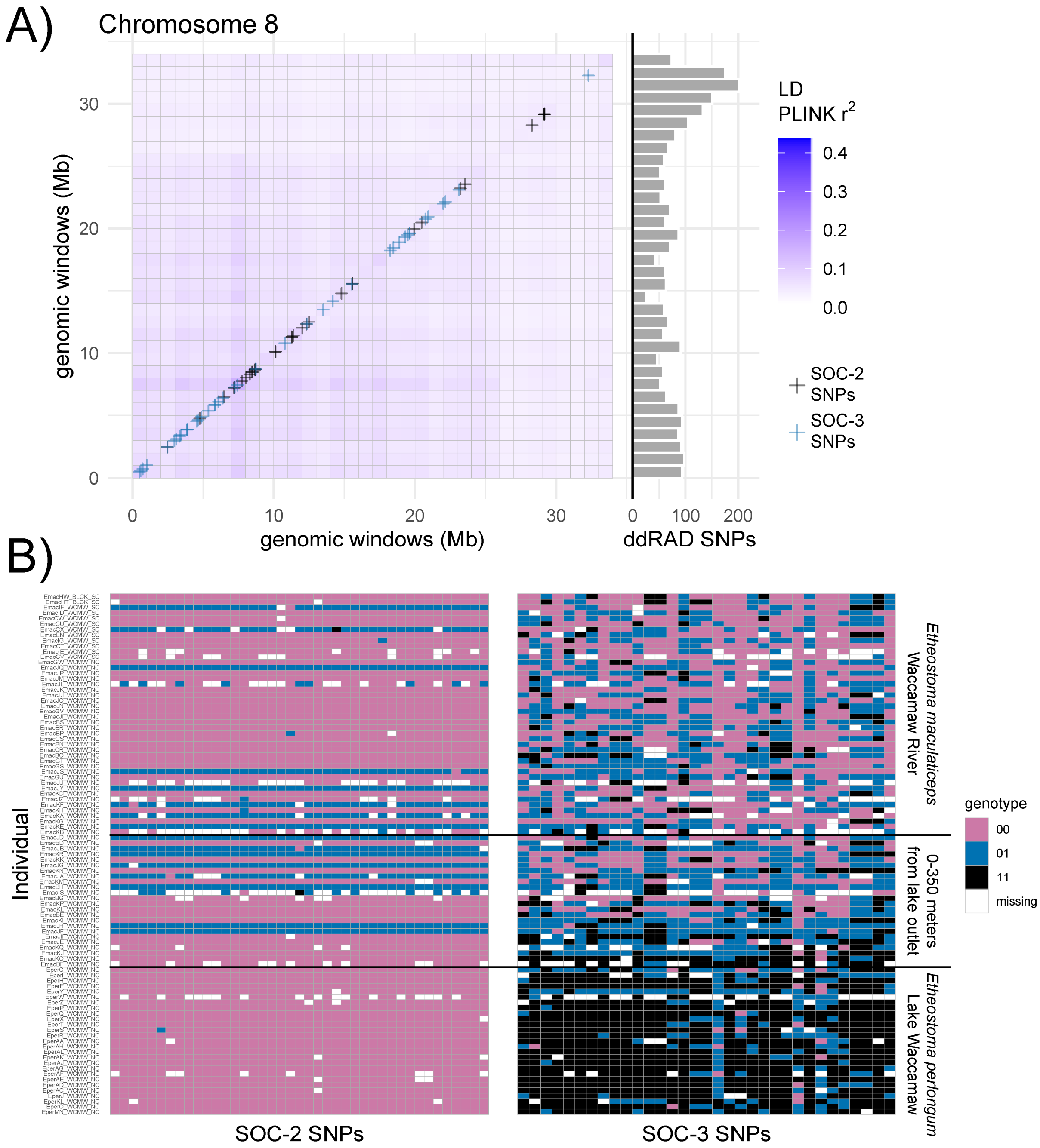

**Fig. S13.** Chromosome 8 outlier clusters of high LD. A) Linkage disequilibrium (LD) estimates for 1 Mbp windows along chromosome 8. Locations of SNPs identified as LD single outlier clusters (SOC-2 and SOC-3) are noted. Histogram shows the distribution of ddRAD SNPs. C) Genotypes of SOC-2 and SOC-3 SNPs. Individuals are ordered by increasing downstream distance from the north shore of Lake Waccamaw.

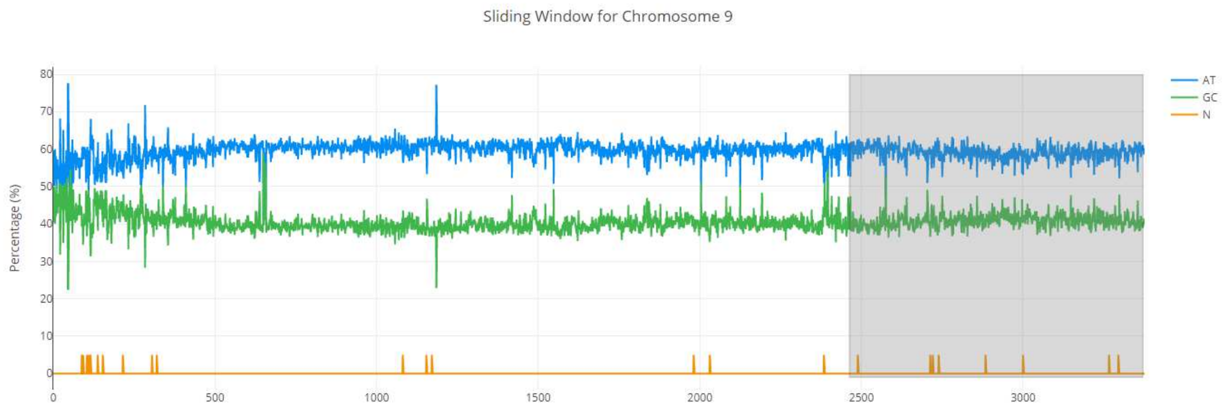

**Fig. S14.** GC content and location of Ns (scaffold break points) along *Etheostoma perlongum* chromosome 9. Gray box indicates the position of the 9 Mb inversion.

**Table S1.** ddRAD samples. YPM = Yale Peabody Museum of Natural History, UT = University of Tennessee, NCSM = North Carolina Museum of Natural Sciences.

| Museum | Catalog Number | Tissue Number | Species | Sample Code | Latitude | Longitude |
|---|---|---|---|---|---|---|
| YPM | ICH.016091 | 8779 | *Etheostoma maculaticeps* | EmacGO_STJH_FL | 29.50937 | -81.946 |
| YPM | ICH.016089 | 8781 | *Etheostoma maculaticeps* | EmacGP_STJH_FL | 29.50937 | -81.946 |
| YPM | ICH.015650 | 8455 | *Etheostoma maculaticeps* | EmacGN_SVNH_GA | 32.1019 | -82.5685 |
| UT | 91.7226 | 6195 | *Etheostoma maculaticeps* | EmacGM_OCGE_GA | 32.49472 | -81.5555 |
| YPM | ICH.030255 | 20274 | *Etheostoma maculaticeps* | EmacGR_OMLG_GA | 32.79933 | -83.8661 |
| YPM | ICH.030396 | 20264 | *Etheostoma maculaticeps* | EmacGQ_OMLG_GA | 33.11392 | -83.8703 |
| UT | 91.7254 | 6094 | *Etheostoma maculaticeps* | EmacGL_SVNH_SC | 33.40692 | -81.7859 |
| YPM | ICH.033027 | 37751 | *Etheostoma maculaticeps* | EmacHT_BLCK_SC | 33.51014 | -79.5582 |
| YPM | ICH.033027 | 37752 | *Etheostoma maculaticeps* | EmacHW_BLCK_SC | 33.51014 | -79.5582 |
| UT | 91.7231 | 6080 | *Etheostoma maculaticeps* | EmacGK_SVNH_SC | 33.91884 | -82.2244 |
| YPM | ICH.029280 | 27175 | *Etheostoma maculaticeps* | EmacGW_WCMW_NC | 33.94111 | -78.6103 |
| YPM | ICH.028323 | 16804 | *Etheostoma maculaticeps* | EmacID_WCMW_SC | 33.95433 | -78.7186 |
| YPM | ICH.028323 | 16805 | *Etheostoma maculaticeps* | EmacIE_WCMW_SC | 33.95433 | -78.7186 |
| YPM | ICH.028323 | 16806 | *Etheostoma maculaticeps* | EmacIF_WCMW_SC | 33.95433 | -78.7186 |
| YPM | ICH.028323 | 16807 | *Etheostoma maculaticeps* | EmacIG_WCMW_SC | 33.95433 | -78.7186 |
| YPM | ICH.028324 | 16809 | *Etheostoma maculaticeps* | EmacCT_WCMW_SC | 33.95433 | -78.7186 |
| YPM | ICH.028325 | 16810 | *Etheostoma maculaticeps* | EmacCU_WCMW_SC | 33.95433 | -78.7186 |
| YPM | ICH.028326 | 16811 | *Etheostoma maculaticeps* | EmacCV_WCMW_SC | 33.95433 | -78.7186 |
| YPM | ICH.028327 | 16812 | *Etheostoma maculaticeps* | EmacCW_WCMW_SC | 33.95433 | -78.7186 |
| YPM | ICH.028328 | 16814 | *Etheostoma maculaticeps* | EmacCX_WCMW_SC | 33.95433 | -78.7186 |
| YPM | ICH.028323 | 16815 | *Etheostoma maculaticeps* | EmacEN_WCMW_SC | 33.95433 | -78.7186 |
| NCSM | 51921 | 51921_1 | *Etheostoma maculaticeps* | EmacBN_WCMW_NC | 34.015 | -78.6316 |
| NCSM | 51921 | 51921_2 | *Etheostoma maculaticeps* | EmacBO_WCMW_NC | 34.015 | -78.6316 |
| NCSM | 51921 | 51921_3 | *Etheostoma maculaticeps* | EmacBP_WCMW_NC | 34.015 | -78.6316 |
| NCSM | 51921 | 51921_4 | *Etheostoma maculaticeps* | EmacBR_WCMW_NC | 34.015 | -78.6316 |
| NCSM | 51921 | 51921_5 | *Etheostoma maculaticeps* | EmacBS_WCMW_NC | 34.015 | -78.6316 |
| NCSM | 51921 | 51921_6 | *Etheostoma maculaticeps* | EmacCR_WCMW_NC | 34.015 | -78.6316 |
| NCSM | 51921 | 51921_7 | *Etheostoma maculaticeps* | EmacCS_WCMW_NC | 34.015 | -78.6316 |
| YPM | ICH.028947 | 27059 | *Etheostoma maculaticeps* | EmacGV_WCMW_NC | 34.04903 | -78.6351 |
| YPM | ICH.028948 | 27060 | *Etheostoma maculaticeps* | EmacJI_WCMW_NC | 34.04903 | -78.6351 |
| YPM | ICH.028949 | 27061 | *Etheostoma maculaticeps* | EmacJJ_WCMW_NC | 34.04903 | -78.6351 |

| | | | | | | |
|---|---|---|---|---|---|---|
| YPM | ICH.028950 | 27062 | *Etheostoma maculaticeps* | EmacJK_WCMW_NC | 34.04903 | -78.6351 |
| YPM | ICH.028951 | 27063 | *Etheostoma maculaticeps* | EmacJL_WCMW_NC | 34.04903 | -78.6351 |
| YPM | ICH.028952 | 27064 | *Etheostoma maculaticeps* | EmacJM_WCMW_NC | 34.04903 | -78.6351 |
| YPM | ICH.028953 | 27065 | *Etheostoma maculaticeps* | EmacJN_WCMW_NC | 34.04903 | -78.6351 |
| YPM | ICH.028954 | 27066 | *Etheostoma maculaticeps* | EmacJO_WCMW_NC | 34.04903 | -78.6351 |
| YPM | ICH.028955 | 27067 | *Etheostoma maculaticeps* | EmacJP_WCMW_NC | 34.04903 | -78.6351 |
| YPM | ICH.028956 | 27068 | *Etheostoma maculaticeps* | EmacJQ_WCMW_NC | 34.04903 | -78.6351 |
| YPM | ICH.029278 | 27023 | *Etheostoma maculaticeps* | EmacGS_WCMW_NC | 34.09456 | -78.5498 |
| YPM | ICH.029278 | 27025 | *Etheostoma maculaticeps* | EmacGT_WCMW_NC | 34.09456 | -78.5498 |
| YPM | ICH.029279 | 27026 | *Etheostoma maculaticeps* | EmacJS_WCMW_NC | 34.09456 | -78.5498 |
| YPM | ICH.029280 | 27028 | *Etheostoma maculaticeps* | EmacJU_WCMW_NC | 34.09456 | -78.5498 |
| YPM | ICH.029278 | 27029 | *Etheostoma maculaticeps* | EmacGU_WCMW_NC | 34.09456 | -78.5498 |
| YPM | ICH.035483 | 26995 | *Etheostoma maculaticeps* | EmacJY_WCMW_NC | 34.158 | -78.5719 |
| YPM | ICH.035483 | 26996 | *Etheostoma maculaticeps* | EmacJZ_WCMW_NC | 34.158 | -78.5719 |
| YPM | ICH.035483 | 26997 | *Etheostoma maculaticeps* | EmacKA_WCMW_NC | 34.158 | -78.5719 |
| YPM | ICH.035483 | 26998 | *Etheostoma maculaticeps* | EmacKB_WCMW_NC | 34.158 | -78.5719 |
| YPM | ICH.035483 | 27000 | *Etheostoma maculaticeps* | EmacKD_WCMW_NC | 34.158 | -78.5719 |
| YPM | ICH.035483 | 27001 | *Etheostoma maculaticeps* | EmacKE_WCMW_NC | 34.158 | -78.5719 |
| YPM | ICH.035483 | 27002 | *Etheostoma maculaticeps* | EmacKF_WCMW_NC | 34.158 | -78.5719 |
| YPM | ICH.035483 | 27003 | *Etheostoma maculaticeps* | EmacKG_WCMW_NC | 34.158 | -78.5719 |
| YPM | ICH.035483 | 27004 | *Etheostoma maculaticeps* | EmacKH_WCMW_NC | 34.158 | -78.5719 |
| YPM | ICH.028437 | 16749 | *Etheostoma maculaticeps* | EmacBD_WCMW_NC | 34.25852 | -78.5232 |
| YPM | ICH.028437 | 16750 | *Etheostoma maculaticeps* | EmacBF_WCMW_NC | 34.25852 | -78.5232 |
| YPM | ICH.028437 | 16751 | *Etheostoma maculaticeps* | EmacBE_WCMW_NC | 34.25852 | -78.5232 |
| YPM | ICH.028437 | 16752 | *Etheostoma maculaticeps* | EmacBG_WCMW_NC | 34.25852 | -78.5232 |
| YPM | ICH.028437 | 16753 | *Etheostoma maculaticeps* | EmacBH_WCMW_NC | 34.25852 | -78.5232 |
| YPM | ICH.029283 | 27156 | *Etheostoma maculaticeps* | EmacKR_WCMW_NC | 34.25852 | -78.5232 |
| YPM | ICH.029283 | 27157 | *Etheostoma maculaticeps* | EmacJA_WCMW_NC | 34.25852 | -78.5232 |
| YPM | ICH.029283 | 27158 | *Etheostoma maculaticeps* | EmacJB_WCMW_NC | 34.25852 | -78.5232 |
| YPM | ICH.029283 | 27160 | *Etheostoma maculaticeps* | EmacJD_WCMW_NC | 34.25852 | -78.5232 |
| YPM | ICH.029283 | 27161 | *Etheostoma maculaticeps* | EmacJE_WCMW_NC | 34.25852 | -78.5232 |
| YPM | ICH.029283 | 27163 | *Etheostoma maculaticeps* | EmacJF_WCMW_NC | 34.25852 | -78.5232 |
| YPM | ICH.029283 | 27164 | *Etheostoma maculaticeps* | EmacJG_WCMW_NC | 34.25852 | -78.5232 |
| YPM | ICH.029283 | 27165 | *Etheostoma maculaticeps* | EmacJH_WCMW_NC | 34.25852 | -78.5232 |

| | | | | | | |
|---|---|---|---|---|---|---|
| YPM | ICH.029283 | 27166 | *Etheostoma maculaticeps* | EmacKI_WCMW_NC | 34.25852 | -78.5232 |
| YPM | ICH.029283 | 27167 | *Etheostoma maculaticeps* | EmacKK_WCMW_NC | 34.25852 | -78.5232 |
| YPM | ICH.029283 | 27168 | *Etheostoma maculaticeps* | EmacKL_WCMW_NC | 34.25852 | -78.5232 |
| YPM | ICH.029283 | 27169 | *Etheostoma maculaticeps* | EmacKM_WCMW_NC | 34.25852 | -78.5232 |
| YPM | ICH.029283 | 27170 | *Etheostoma maculaticeps* | EmacKN_WCMW_NC | 34.25852 | -78.5232 |
| YPM | ICH.029283 | 27171 | *Etheostoma maculaticeps* | EmacKO_WCMW_NC | 34.25852 | -78.5232 |
| YPM | ICH.029283 | 27172 | *Etheostoma maculaticeps* | EmacKP_WCMW_NC | 34.25852 | -78.5232 |
| YPM | ICH.029283 | 27173 | *Etheostoma maculaticeps* | EmacKQ_WCMW_NC | 34.25852 | -78.5232 |
| YPM | ICH.029283 | 27174 | *Etheostoma maculaticeps* | EmacKJ_WCMW_NC | 34.25852 | -78.5232 |
| YPM | ICH.031397 | 31123 | *Etheostoma maculaticeps* | EmacIS_WCMW_NC | 34.25865 | -78.5232 |
| YPM | ICH.031397 | 31124 | *Etheostoma maculaticeps* | EmacII_WCMW_NC | 34.25865 | -78.5232 |
| UT | 91.7228 | 6013 | *Etheostoma maculaticeps* | EmacGI_SNTE_SC | 34.26506 | -81.6633 |
| UT | 91.7228 | 6018 | *Etheostoma maculaticeps* | EmacGJ_SNTE_SC | 34.26508 | -81.6632 |
| NCSM | 33204 | 33204_1 | *Etheostoma maculaticeps* | EmacBI_PEDE_NC | 34.388 | -79.0012 |
| NCSM | 33204 | 33204_2 | *Etheostoma maculaticeps* | EmacBJ_PEDE_NC | 34.388 | -79.0012 |
| NCSM | 33204 | 33204_3 | *Etheostoma maculaticeps* | EmacBK_PEDE_NC | 34.388 | -79.0012 |
| YPM | ICH.032682 | 37320 | *Etheostoma maculaticeps* | EmacHM_LUMB_NC | 34.68301 | -79.3763 |
| NCSM | 53551 | 53551_1 | *Etheostoma maculaticeps* | EmacAQ_FEAR_NC | 34.7222 | -77.9817 |
| NCSM | 53551 | 53551_2 | *Etheostoma maculaticeps* | EmacAR_FEAR_NC | 34.7222 | -77.9817 |
| YPM | ICH.032696 | 37322 | *Etheostoma maculaticeps* | EmacHO_LUMB_NC | 34.88364 | -79.4517 |
| YPM | ICH.032750 | 37313 | *Etheostoma maculaticeps* | EmacHK_LUMB_NC | 34.97511 | -79.3582 |
| YPM | ICH.032750 | 37314 | *Etheostoma maculaticeps* | EmacHL_LUMB_NC | 34.97511 | -79.3582 |
| NCSM | 29535 | 29535_1 | *Etheostoma maculaticeps* | EmacBL_PEDE_NC | 35.0886 | -79.8354 |
| NCSM | 29535 | 29535_2 | *Etheostoma maculaticeps* | EmacBM_PEDE_NC | 35.0886 | -79.8354 |
| YPM | ICH.032790 | 37260 | *Etheostoma maculaticeps* | EmacHI_LUMB_NC | 35.1342 | -79.4288 |
| YPM | ICH.032790 | 37261 | *Etheostoma maculaticeps* | EmacHJ_LUMB_NC | 35.1342 | -79.4288 |
| YPM | ICH.033033 | 37740 | *Etheostoma maculaticeps* | EmacHR_LUMB_NC | 35.20782 | -79.6648 |
| YPM | ICH.033033 | 37741 | *Etheostoma maculaticeps* | EmacHS_LUMB_NC | 35.20782 | -79.6648 |
| YPM | ICH.033035 | 37728 | *Etheostoma maculaticeps* | EmacHP_LUMB_NC | 35.2699 | -79.6786 |
| YPM | ICH.033035 | 37729 | *Etheostoma maculaticeps* | EmacHQ_LUMB_NC | 35.2699 | -79.6786 |
| NCSM | 61004 | 61004_2 | *Etheostoma maculaticeps* | EmacEU_FEAR_NC | 35.5145 | -78.7861 |
| NCSM | 61004 | 61004_3 | *Etheostoma maculaticeps* | EmacET_FEAR_NC | 35.5145 | -78.7861 |
| NCSM | 30566 | 30566_1 | *Etheostoma maculaticeps* | EmacFG_FEAR_NC | 36.2281 | -79.3735 |
| YPM | ICH.031024 | 29812 | *Etheostoma maculaticeps* | EmacGY_NEW_NC | 36.47667 | -81.118 |

| | | | | | | |
|---|---|---|---|---|---|---|
| YPM | ICH.031024 | 29813 | *Etheostoma maculaticeps* | EmacGZ_NEW_NC | 36.47667 | -81.118 |
| YPM | ICH.031024 | 29814 | *Etheostoma maculaticeps* | EmacHA_NEW_NC | 36.47667 | -81.118 |
| YPM | ICH.031024 | 29815 | *Etheostoma maculaticeps* | EmacHB_NEW_NC | 36.47667 | -81.118 |
| YPM | ICH.031024 | 29816 | *Etheostoma maculaticeps* | EmacHC_NEW_NC | 36.47667 | -81.118 |
| NCSM | 53328 | 53328_1 | *Etheostoma maculaticeps* | EmacGX_NEW_NC | 36.49969 | -81.0364 |
| YPM | ICH.031057 | 29841 | *Etheostoma maculaticeps* | EmacHD_PEDE_NC | 36.51061 | -80.5915 |
| YPM | ICH.031057 | 29843 | *Etheostoma maculaticeps* | EmacHE_PEDE_NC | 36.51061 | -80.5915 |
| YPM | ICH.031057 | 29844 | *Etheostoma maculaticeps* | EmacHF_PEDE_NC | 36.51061 | -80.5915 |
| YPM | ICH.031057 | 29845 | *Etheostoma maculaticeps* | EmacHG_PEDE_NC | 36.51061 | -80.5915 |
| YPM | ICH.031057 | 29846 | *Etheostoma maculaticeps* | EmacHH_PEDE_NC | 36.51061 | -80.5915 |
| YPM | ICH.030277 | 20373 | *Etheostoma nigrum* | EngrDL_OBON_TN | 36.27772 | -88.3875 |
| YPM | ICH.027398 | 24567 | *Etheostoma nigrum* | EngrEX_CUMB_KY | 37.0756 | -82.7721 |
| NCSM | 74833 | 74833_1 | *Etheostoma olmstedi* | EolmCO_CSPK_VA | 37.8771 | -76.4949 |
| YPM | ICH.030863 | 17450 | *Etheostoma olmstedi* | EolmNH_OSWG_NY | 43.01407 | -76.3371 |
| YPM | ICH.034479 | 26967 | *Etheostoma perlongum* | EperP_WCMW_NC | 34.25967 | -78.4768 |
| YPM | ICH.034479 | 26968 | *Etheostoma perlongum* | EperQ_WCMW_NC | 34.25967 | -78.4768 |
| YPM | ICH.034480 | 26969 | *Etheostoma perlongum* | EperR_WCMW_NC | 34.25967 | -78.4768 |
| YPM | ICH.034481 | 26971 | *Etheostoma perlongum* | EperS_WCMW_NC | 34.25967 | -78.4768 |
| YPM | ICH.034482 | 26973 | *Etheostoma perlongum* | EperT_WCMW_NC | 34.25967 | -78.4768 |
| YPM | ICH.034483 | 26976 | *Etheostoma perlongum* | EperW_WCMW_NC | 34.25967 | -78.4768 |
| YPM | ICH.034484 | 26977 | *Etheostoma perlongum* | EperX_WCMW_NC | 34.25967 | -78.4768 |
| YPM | ICH.034485 | 26978 | *Etheostoma perlongum* | EperY_WCMW_NC | 34.25967 | -78.4768 |
| YPM | ICH.034486 | 26979 | *Etheostoma perlongum* | EperZ_WCMW_NC | 34.25967 | -78.4768 |
| YPM | ICH.034487 | 26980 | *Etheostoma perlongum* | EperAA_WCMW_NC | 34.25967 | -78.4768 |
| NCSM | 51791 | 51791_1 | *Etheostoma perlongum* | EperG_WCMW_NC | 34.2611 | -78.5223 |
| NCSM | 51791 | 51791_2 | *Etheostoma perlongum* | EperH_WCMW_NC | 34.2611 | -78.5223 |
| NCSM | 51791 | 51791_3 | *Etheostoma perlongum* | EperI_WCMW_NC | 34.2611 | -78.5223 |
| YPM | ICH.028448 | 16736 | *Etheostoma perlongum* | EperE_WCMW_NC | 34.2611 | -78.5223 |
| NCSM | 51798 | 51798_1 | *Etheostoma perlongum* | EperJ_WCMW_NC | 34.2904 | -78.4745 |
| NCSM | 51798 | 51798_2 | *Etheostoma perlongum* | EperKL_WCMW_NC | 34.2904 | -78.4745 |
| YPM | ICH.032699 | 37173 | *Etheostoma perlongum* | EperAG_WCMW_NC | 34.30049 | -78.5514 |
| YPM | ICH.032699 | 37175 | *Etheostoma perlongum* | EperAH_WCMW_NC | 34.30049 | -78.5514 |
| YPM | ICH.032699 | 37255 | *Etheostoma perlongum* | EperAC_WCMW_NC | 34.30049 | -78.5514 |
| YPM | ICH.032699 | 37256 | *Etheostoma perlongum* | EperAD_WCMW_NC | 34.30049 | -78.5514 |

| | | | | | | |
|---|---|---|---|---|---|---|
| YPM | ICH.032699 | 37257 | *Etheostoma perlongum* | EperAE_WCMW_NC | 34.30049 | -78.5514 |
| YPM | ICH.032699 | 37258 | *Etheostoma perlongum* | EperAF_WCMW_NC | 34.30049 | -78.5514 |
| YPM | ICH.033096 | 37806 | *Etheostoma perlongum* | EperAJ_WCMW_NC | 34.30049 | -78.5514 |
| YPM | ICH.033096 | 37810 | *Etheostoma perlongum* | EperAK_WCMW_NC | 34.30049 | -78.5514 |
| YPM | ICH.033096 | 37811 | *Etheostoma perlongum* | EperAL_WCMW_NC | 34.30049 | -78.5514 |
| NCSM | 56309 | 56309_1 | *Etheostoma perlongum* | EperMN_WCMW_NC | 34.3166 | -78.5243 |
| NCSM | 56309 | 56309_2 | *Etheostoma perlongum* | EperO_WCMW_NC | 34.3166 | -78.5243 |

**Table S2.** iPyrad assembly parameters. See https://ipyrad.readthedocs.io/ for more information about each parameter.

| IPYRAD ASSEMBLY PARAMTER | SETTING |
|---|---|
| ASSEMBLY_METHOD | reference |
| DATATYPE | ddrad |
| RESTRICTION_OVERHANG | TGCAG, CCG |
| MAX_LOW_QUAL_BASES | 5 |
| PHRED_QSCORE_OFFSET | 33 |
| MINDEPTH_STATISTICAL | 6 |
| MINDEPTH_MAJRULE | 6 |
| MAXDEPTH | 10,000 |
| MAX_BARCODE_MISMATCH | 0 |
| FILTER_ADAPTERS | 2 |
| FILTER_MIN_TRIM_LEN | 35 |
| MAX_ALLELES_CONSENS | 2 |
| MAX_NS_CONSENS | 0.05 |
| MAX_HS_CONSENS | 0.05 |
| MAX_SNPS_LOCUS | 0.2 |
| MAX_INDELS_LOCUS | 8 |
| MAX_SHARED_HS_LOCUS | 0.5 |
| TRIM_READS | 0, 0, 0, 0 |
| TRIM_LOCI | 0, 0, 0, 0 |
| MIN_SAMPLES_LOCUS | 98 (30% missing data)<br>112 (20% missing data)<br>126 (10% missing data)<br>133 (5% missing data) |

**Table S3.** *Etheostoma perlongum* genome assembly statistics.

| | | |
|---|---|---|
| **Assembly** | **Number of contigs (initial assembly)** | **903** |
| | Longest contig (initial assembly) (Mb) | 9.9 |
| | Contig N50 (initial assembly) (Mb) | 2.5 |
| | Final assembly length (Mb) | 788.7 |
| | Number of scaffolds (chromosomes) | 1,179 |
| | Number of contigs (final assembly) | 2,095 |
| | Contig N50 (final assembly) (Mb) | 2.3 |
| | Scaffold N50 (Mb) | 30.8 |
| | GC (%) | 40.5 |
| **BUSCO v5 (actinopterygii_odb10)** | Complete (%) | 96.7 |
| | Complete single copy (%) | 95.7 |
| | Complete duplicated (%) | 1 |
| | Fragmented (%) | 1 |
| | Missing (%) | 2.3 |
| **Annotation** | Predicted genes | 28,034 |
| | Mean [median] gene length (bp) | 11,370.6 [7,482] |
| | Mean [median] exon length (bp) | 204.1 [126] |
| | Mean [median] intron length (bp) | 1,150.8 [482] |
| | Mean [median] exons per gene | 9.2 [6] |
| | Mean [median] introns per gene | 8.2 [5] |

**Table S4.** ddRAD assembly statistics for phylogenetic analyses.

| ***Maximum percent missing samples per locus*** | ***Loci*** | ***Variable sites*** | ***Parsimony informative sites*** | ***Alignment length (bp)*** | ***Percent missing sites in alignment*** |
|---|---|---|---|---|---|
| *30%* | 66,126 | 510,618 | 263,539 | 9,127,827 | 39% |
| *20%* | 46,874 | 371,480 | 192,551 | 6,725,088 | 37% |
| *10%* | 25,031 | 199,073 | 103,788 | 3,685,299 | 34% |
| *5%* | 11,440 | 90,498 | 47,014 | 1,711,662 | 33% |

**Table S5.** Fastsimcoal2 ΔAIC table. Best fit model indicated by black shading, models with ΔAIC less than 10 indicated by dark gray shading.

| LAKE-STREAM DIVERGENCE (GENERATIONS) | MODEL 1 ΔAIC | MODEL 2 ΔAIC | MODEL 3 ΔAIC |
|---|---|---|---|
| **1,000** | 2293.5 | 76.1 | 100.5 |
| **2,000** | 2287.9 | 28.6 | 158.3 |
| **3,000** | 2236.1 | 194.6 | 159.2 |
| **4,000** | 2274.5 | 219.1 | 19.8 |
| **5,000** | 2385.8 | 81.8 | 26.2 |
| **6,000** | 2565.3 | 255.9 | 10.8 |
| **7,000** | 2806.4 | 12.2 | 19.8 |
| **8,000** | 3180.0 | 208.6 | 19.5 |
| **9,000** | 3483.7 | 17.5 | 29.7 |
| **10,000** | 3860.9 | 25.6 | 15.2 |
| **11,000** | 4203.1 | 0.0 | 32.5 |
| **12,000** | 4695.7 | 18.1 | 11.4 |
| **13,000** | 5158.9 | 17.2 | 10.7 |
| **14,000** | 5558.9 | 57.0 | 4.8 |
| **15,000** | 6031.4 | 33.9 | 10.0 |
| **16,000** | 6430.4 | 24.0 | 10.4 |
| **17,000** | 6935.9 | 228.7 | 12.9 |
| **18,000** | 7247.9 | 18.1 | 23.3 |
| **19,000** | 7691.1 | 66.5 | 33.4 |
| **20,000** | 8090.9 | 19.3 | 25.1 |
| **21,000** | 8572.8 | 28.8 | 43.6 |
| **22,000** | 8919.8 | 44.7 | 38.8 |
| **23,000** | 9313.7 | 37.5 | 30.0 |
| **24,000** | 9596.3 | 102.6 | 77.6 |
| **25,000** | 10015.2 | 197.6 | 31.0 |
| **26,000** | 10312.0 | 68.2 | 56.0 |
| **27,000** | 10769.9 | 73.1 | 52.4 |
| **28,000** | 11138.7 | 246.3 | 36.1 |
| **29,000** | 11459.3 | 84.2 | 88.5 |
| **30,000** | 11860.5 | 97.6 | 70.3 |
| **31,000** | 12187.2 | 99.2 | 156.6 |
| **32,000** | 12385.0 | 95.1 | 169.0 |

**Table S6.** Vision, olfaction, and circadian rhythm genes located within the Chr9 inversion.

| *E. perlongum* gene ID | Inversion $D_{XY}$ peak | *D. rerio* ortholog | Functional Significance |
|---|---|---|---|
| **Eper_00008530**[*] | 1 | *barhl2* | Regulates retinal cell differentiation during development (Schuhmacher et al. 2011). |
| **Eper_00008535**[+] | 1 | *lmo4b* | Transcription factor, negative regulator of forebrain size (olfaction-related) and is expressed in the optic primordia (McCollum et al. 2007). Regulates *six3*, a negative regulator of *bmp4* signaling controlling craniofacial development (Gestri et al. 2005). |
| **Eper_00008679**[^] | none | *rorca* | Activated by CLOCK:ARNTL dimer, mutations in *rorca* disrupt diurnal regulation and activity in cavefish (Mack et al. 2021). |
| **Eper_00008692** | 2 | *pde6d* | Encodes for a key enzyme in the phototransduction pathway, a retinal rod rhodopsin-sensitive cGMP phosphodiesterase (Karan et al. 2008; Ershova et al. 1997). Plays a role in retinal dystrophy in Joubert Syndrome 22 (Thomas et al. 2014). |
| **Eper_00008726**[^] | 2 | *prkaa2* | Links nutrient availability and energy metabolism to circadian oscillations (Vieira et al. 2008, Lamia et al. 2009) |
| **Eper_00008708**[*] | 2 | *prdx6* | Implicated in bullous retinoschisis, a disease involving separation of retinal cell layers (Sudha et al. 2018). |
| **Eper_00008758**[*] | 2 | *nrl* | Master regulator of retinal development, specifically rod photoreceptor differentiation; regulates rhodopsin and short- and long-wave opsin expression (Oel et al. 2020). |
| **Eper_00008768**[^] | 2 | *clock1a* | Core circadian rhythm transcription factor that controls circadian rhythmicity and its product forms a heterodimer with BMAL1 to act as an upstream regulator of transcription of thousands of genes (Rey et al. 2011). Implicated in reproductive seasonality in fishes (Krabbenhoft and Turner 2014). |
| **Eper_00008772**[*] | 2 | *kita* | Regulated by gonadotropins and crucial for gametogenesis in catfish (Laldinsangi and Senthilkumaran 2018), involved in melanocyte development and differentiation (Mellgren and Johnson 2004) |
| **Eper_00008774**[^] | 2 | *gsx2* | Homeobox transcription factor that regulates brain development, particularly olfactory bulb interneurons (Coltogirone et al. 2021). |
| **Eper_00008808**[^] | none | *prdm5* | Mutations in prdm5 cause brittle cornea |

| | | | |
|---|---|---|---|
| | | | syndrome in humans (Wright et al. 2011) |
| **Eper_00008815 Eper_00008817** | none | *reps2* | A microdeletion in *reps2* is associated with Nance-Horan Syndrome (Liao et al. 2011). |
| **Eper_00008818** | none | *nhsa* | Involved in Nance-Horan syndrome, which involves vision loss and microphthalmia, and X-linked cataract-40 (Burdon et al. 2003; Sharma et al. 2008). |
| **Eper_00008825** | none | *rs1a* | Target of *prdx6* (Sudha et al. 2018). Secretory protein expressed exclusively in the retina that plays a crucial role in cellular organization; mutations cause retinoschisis, a disease involving splitting of the retina's neurosensory layers, leading to vision loss (Gehrig et al.1999). |
| **Eper_00008828 Eper_00008829[+]** | none | *shroom2* | Key regulator of retinal pigment epithelium (RPE) pigmentation and suspected of contributing to ocular albinism, which disrupts normal vision (Fairbank et al. 2006). |
| **Eper_00008856[+]** | none | *oca2* | Produces the P protein in melanocytes, including in the retinal pigment epithelium. Mutations in this gene cause type 2 oculocutaneous albinism (Stevens et al. 1997) |
| **Eper_00008872[+]** | none | *dpt* | Expressed in brain and optic nerve cells, along with four layers of the retina responsible for optic nerve transmission (i.e., ganglion cell layer, inner nuclear layer, outer plexiform layer, and outer nuclear layer (Stenkamp et al. 2007, Tan et al. 2013). |

[*]syntenic with *Gadus morhua* LG12 supergene and *Oncorhynchus mykiss* Omy5 supergene, [+]syntenic with *Gadus morhua* LG12 supergene, [^]syntenic with *Oncorhynchus mykiss* Omy5 supergene.

**Table S7.** Enriched GO terms retained by REVIGO after semantic similarity clustering for the chromosome $F_{ST}$ plateau, Dxy peak 1, and Dxy peak 2.

| GO term ID | description | frequency | uniqueness | dispensability |
|---|---|---|---|---|
| | **$F_{ST}$ plateau** | | | |
| **GO:0001952** | “regulation of cell-matrix adhesion" | 0.02 | 0.88 | 0 |
| **GO:0015693** | “magnesium ion transport" | 0.107 | 0.953 | 0 |
| **GO:0032502** | “developmental process" | 2.266 | 1 | 0 |
| **GO:0048730** | “epidermis morphogenesis" | 0.006 | 0.916 | 0 |
| **GO:0070293** | “renal absorption" | 0.003 | 0.929 | 0 |
| **GO:0071391** | “cellular response to estrogen stimulus" | 0.006 | 0.933 | 0 |
| **GO:0090161** | “Golgi ribbon formation" | 0.003 | 0.927 | 0.005 |
| **GO:0006687** | “glycosphingolipid metabolic process" | 0.02 | 0.932 | 0.005 |
| **GO:0035249** | “synaptic transmission glutamatergic" | 0.006 | 0.901 | 0.005 |
| **GO:0070979** | “protein K11-linked ubiquitination" | 0.008 | 0.923 | 0.085 |
| **GO:0070189** | “kynurenine metabolic process" | 0.098 | 0.928 | 0.099 |
| **GO:0098901** | “regulation of cardiac muscle cell action potential" | 0.004 | 0.856 | 0.131 |
| **GO:0010874** | “regulation of cholesterol efflux" | 0.006 | 0.872 | 0.134 |
| **GO:0032485** | “regulation of Ral protein signal transduction" | 0.002 | 0.834 | 0.135 |
| **GO:0045682** | “regulation of epidermis development" | 0.011 | 0.866 | 0.139 |
| **GO:0042795** | “snRNA transcription by RNA polymerase II" | 0.004 | 0.913 | 0.139 |
| **GO:0060968** | “regulation of gene silencing" | 0.027 | 0.875 | 0.155 |
| **GO:0006515** | “protein quality control for misfolded or incompletely synthesized proteins" | 0.062 | 0.886 | 0.19 |
| **GO:0051569** | “regulation of histone H3-K4 methylation" | 0.009 | 0.872 | 0.203 |
| **GO:0016192** | “vesicle-mediated transport" | 1.371 | 0.945 | 0.242 |
| **GO:0000350** | “generation of catalytic spliceosome for second transesterification step" | 0.008 | 0.879 | 0.244 |
| **GO:0070059** | “intrinsic apoptotic signaling pathway in response to endoplasmic reticulum stress" | 0.006 | 0.823 | 0.251 |
| **GO:0018342** | “protein prenylation" | 0.038 | 0.923 | 0.253 |
| **GO:0000122** | “negative regulation of transcription by RNA polymerase II" | 0.189 | 0.831 | 0.259 |
| **GO:0008612** | “peptidyl-lysine modification to peptidyl-hypusine" | 0.02 | 0.9 | 0.268 |
| **GO:0006233** | “dTDP biosynthetic process" | 0.038 | 0.893 | 0.299 |
| **GO:0051649** | “establishment of localization in cell" | 1.968 | 0.908 | 0.327 |
| **GO:0007098** | “centrosome cycle" | 0.031 | 0.906 | 0.328 |
| **GO:0007186** | “G protein-coupled receptor signaling pathway" | 1.081 | 0.78 | 0.346 |
| **GO:0006695** | “cholesterol biosynthetic process" | 0.014 | 0.916 | 0.348 |
| **GO:0030104** | “water homeostasis" | 0.011 | 0.843 | 0.354 |
| **GO:0016188** | “synaptic vesicle maturation" | 0.003 | 0.87 | 0.369 |
| **GO:1905455** | “positive regulation of myeloid progenitor cell differentiation" | 0 | 0.857 | 0.378 |
| **GO:0030223** | “neutrophil differentiation" | 0.003 | 0.897 | 0.379 |
| **GO:0001505** | “regulation of neurotransmitter levels" | 0.06 | 0.84 | 0.413 |
| **GO:0006821** | “chloride transport" | 0.139 | 0.953 | 0.421 |

| GO term ID | description | frequency | uniqueness | dispensability |
|---|---|---|---|---|
| **GO:0009143** | “nucleoside triphosphate catabolic process" | 0.112 | 0.872 | 0.434 |
| **GO:0035522** | “monoubiquitinated histone H2A deubiquitination" | 0.003 | 0.86 | 0.438 |
| **GO:0007169** | “transmembrane receptor protein tyrosine kinase signaling pathway" | 0.14 | 0.771 | 0.443 |
| **GO:2000050** | “regulation of non-canonical Wnt signaling pathway" | 0.004 | 0.831 | 0.448 |
| **GO:0060729** | “intestinal epithelial structure maintenance" | 0.002 | 0.801 | 0.453 |
| **GO:0007422** | “peripheral nervous system development" | 0.019 | 0.897 | 0.458 |
| **GO:0090024** | “negative regulation of neutrophil chemotaxis" | 0.001 | 0.805 | 0.46 |
| **GO:1905897** | “regulation of response to endoplasmic reticulum stress" | 0.02 | 0.816 | 0.469 |
| **GO:0048702** | “embryonic neurocranium morphogenesis" | 0.003 | 0.894 | 0.469 |
| **GO:0031580** | “membrane raft distribution" | 0.001 | 0.892 | 0.471 |
| **GO:0045747** | “positive regulation of Notch signaling pathway" | 0.009 | 0.813 | 0.49 |
| **GO:0005576** | “extracellular region" | 2.581 | 1 | 0 |
| **GO:0031462** | “Cul2-RING ubiquitin ligase complex" | 0.009 | 0.719 | 0 |
| **GO:0032391** | “photoreceptor connecting cilium" | 0.009 | 0.683 | 0 |
| **GO:0016020** | “membrane" | 63.598 | 1 | 0 |
| **GO:0099513** | “polymeric cytoskeletal fiber" | 0.721 | 0.656 | 0.142 |
| **GO:0017054** | “negative cofactor 2 complex" | 0.007 | 0.702 | 0.197 |
| **GO:0005762** | “mitochondrial large ribosomal subunit" | 0.054 | 0.624 | 0.359 |
| **GO:0070776** | “MOZ/MORF histone acetyltransferase complex" | 0.008 | 0.59 | 0.445 |
| **GO:0005968** | “Rab-protein geranylgeranyltransferase complex" | 0.018 | 0.687 | 0.465 |
| **GO:0003860** | “3-hydroxyisobutyryl-CoA hydrolase activity" | 0.022 | 0.96 | 0 |
| **GO:0004890** | “GABA-A receptor activity" | 0.017 | 0.881 | 0 |
| **GO:0008073** | “ornithine decarboxylase inhibitor activity" | 0.005 | 0.943 | 0 |
| **GO:0015095** | “magnesium ion transmembrane transporter activity" | 0.088 | 0.959 | 0 |
| **GO:0050661** | “NADP binding" | 0.66 | 0.971 | 0 |
| **GO:0008489** | “UDP-galactose:glucosylceramide beta-1 4-galactosyltransferase activity" | 0 | 0.913 | 0.016 |
| **GO:0000253** | “3-keto sterol reductase activity" | 0.001 | 0.931 | 0.017 |
| **GO:0004639** | “phosphoribosylaminoimidazolesuccinocarboxamide synthase activity" | 0.034 | 0.948 | 0.021 |
| **GO:0016787** | “hydrolase activity" | 19.406 | 0.959 | 0.035 |
| **GO:0017147** | “Wnt-protein binding" | 0.009 | 0.947 | 0.038 |
| **GO:0003730** | “mRNA 3'-UTR binding" | 0.025 | 0.969 | 0.086 |
| **GO:0036137** | “kynurenine aminotransferase activity" | 0.004 | 0.909 | 0.106 |
| **GO:0008146** | “sulfotransferase activity" | 0.116 | 0.899 | 0.141 |
| **GO:0008318** | “protein prenyltransferase activity" | 0.029 | 0.833 | 0.16 |
| **GO:0003865** | “3-oxo-5-alpha-steroid 4-dehydrogenase activity" | 0.008 | 0.924 | 0.169 |
| **GO:0004798** | “thymidylate kinase activity" | 0.032 | 0.879 | 0.17 |
| **GO:0047429** | “nucleoside-triphosphate diphosphatase activity" | 0.138 | 0.956 | 0.176 |
| **GO:0008942** | “nitrite reductase [NAD(P)H] activity" | 0.027 | 0.919 | 0.2 |
| **GO:0019789** | “SUMO transferase activity" | 0.018 | 0.8 | 0.241 |
| **GO:0042922** | “neuromedin U receptor binding" | 0.001 | 0.942 | 0.28 |

| GO term ID | description | frequency | uniqueness | dispensability |
|---|---|---|---|---|
| **GO:0016705** | “oxidoreductase activity acting on paired donors with incorporation or reduction of molecular oxygen" | 1.23 | 0.899 | 0.281 |
| **GO:0050660** | “flavin adenine dinucleotide binding" | 1.629 | 0.969 | 0.282 |
| **GO:0140326** | “ATPase-coupled intramembrane lipid transporter activity" | 0.049 | 0.883 | 0.285 |
| **GO:0016712** | “oxidoreductase activity acting on paired donors with incorporation or reduction of molecular oxygen reduced flavin or flavoprotein as one donor" | 0.039 | 0.885 | 0.289 |
| **GO:0044325** | “ion channel binding" | 0.017 | 0.946 | 0.332 |
| **GO:0008097** | “5S rRNA binding" | 0.04 | 0.969 | 0.336 |
| **GO:0004044** | “amidophosphoribosyltransferase activity" | 0.031 | 0.895 | 0.355 |
| **GO:0005018** | “platelet-derived growth factor alpha-receptor activity" | 0.001 | 0.739 | 0.422 |
| **GO:0004176** | “ATP-dependent peptidase activity" | 0.199 | 0.829 | 0.424 |
| **GO:0004969** | “histamine receptor activity" | 0.003 | 0.869 | 0.457 |
| | **Dxy peak 1** | | | |
| **GO:0060968** | “regulation of gene silencing" | 0.027 | 0.964 | 0 |
| **GO:0061053** | “somite development" | 0.024 | 0.754 | 0 |
| **GO:0070189** | “kynurenine metabolic process" | 0.098 | 0.861 | 0 |
| **GO:0009058** | “biosynthetic process" | 24.708 | 0.965 | 0.042 |
| **GO:0070050** | “neuron cellular homeostasis" | 0.006 | 0.959 | 0.127 |
| **GO:0006520** | “cellular amino acid metabolic process" | 5.778 | 0.85 | 0.45 |
| **GO:0060219** | “camera-type eye photoreceptor cell differentiation" | 0.007 | 0.74 | 0.45 |
| **GO:0030054** | “cell junction" | 0.875 | 1 | 0 |
| **GO:0005923** | “bicellular tight junction" | 0.074 | 1 | 0 |
| **GO:0005198** | “structural molecule activity" | 2.916 | 1 | 0 |
| **GO:0030170** | “pyridoxal phosphate binding" | 1.159 | 1 | 0 |
| **GO:0036137** | “kynurenine aminotransferase activity" | 0.004 | 0.908 | 0 |
| **GO:0008146** | “sulfotransferase activity" | 0.116 | 0.908 | 0.141 |
| | **Dxy peak 2** | | | |
| **GO:0006940** | “regulation of smooth muscle contraction" | 0.012 | 0.902 | 0 |
| **GO:0009648** | “photoperiodism" | 0.009 | 0.926 | 0 |
| **GO:0016051** | “carbohydrate biosynthetic process" | 1.22 | 0.913 | 0 |
| **GO:0048512** | “circadian behavior" | 0.007 | 0.961 | 0 |
| **GO:0050953** | “sensory perception of light stimulus" | 0.076 | 0.916 | 0 |
| **GO:0090161** | “Golgi ribbon formation" | 0.003 | 0.943 | 0 |
| **GO:0060294** | “cilium movement involved in cell motility" | 0.036 | 0.942 | 0.005 |
| **GO:0018149** | “peptide cross-linking" | 0.016 | 0.884 | 0.053 |
| **GO:0006644** | “phospholipid metabolic process" | 1.406 | 0.87 | 0.08 |
| **GO:0001505** | “regulation of neurotransmitter levels" | 0.06 | 0.899 | 0.139 |
| **GO:0051492** | “regulation of stress fiber assembly" | 0.017 | 0.895 | 0.142 |
| **GO:0045746** | “negative regulation of Notch signaling pathway" | 0.009 | 0.857 | 0.144 |

| GO term ID | description | frequency | uniqueness | dispensability |
|---|---|---|---|---|
| **GO:0001952** | “regulation of cell-matrix adhesion" | 0.02 | 0.895 | 0.151 |
| **GO:0030335** | “positive regulation of cell migration" | 0.089 | 0.847 | 0.167 |
| **GO:0030163** | “protein catabolic process" | 1.027 | 0.855 | 0.195 |
| **GO:0009113** | “purine nucleobase biosynthetic process" | 0.123 | 0.842 | 0.199 |
| **GO:0000350** | “generation of catalytic spliceosome for second transesterification step" | 0.008 | 0.875 | 0.244 |
| **GO:0008612** | “peptidyl-lysine modification to peptidyl-hypusine" | 0.02 | 0.863 | 0.255 |
| **GO:0018342** | “protein prenylation" | 0.038 | 0.848 | 0.268 |
| **GO:0035522** | “monoubiquitinated histone H2A deubiquitination" | 0.003 | 0.843 | 0.29 |
| **GO:0006487** | “protein N-linked glycosylation" | 0.116 | 0.815 | 0.301 |
| **GO:1905457** | “negative regulation of lymphoid progenitor cell differentiation" | 0.001 | 0.845 | 0.35 |
| **GO:0001649** | “osteoblast differentiation" | 0.017 | 0.91 | 0.449 |
| **GO:0038093** | “Fc receptor signaling pathway" | 0.004 | 0.791 | 0.45 |
| **GO:0009143** | “nucleoside triphosphate catabolic process" | 0.112 | 0.824 | 0.484 |
| **GO:0000785** | “chromatin" | 0.55 | 0.682 | 0 |
| **GO:0005968** | “Rab-protein geranylgeranyltransferase complex" | 0.018 | 0.563 | 0.082 |
| **GO:0001534** | “radial spoke" | 0.006 | 0.59 | 0.298 |
| **GO:0070776** | “MOZ/MORF histone acetyltransferase complex" | 0.008 | 0.537 | 0.464 |
| **GO:0004639** | “phosphoribosylaminoimidazolesuccinocarboxamide synthase activity" | 0.034 | 0.97 | 0 |
| **GO:0004969** | “histamine receptor activity" | 0.003 | 0.854 | 0 |
| **GO:0050661** | “NADP binding" | 0.66 | 0.967 | 0 |
| **GO:0003865** | “3-oxo-5-alpha-steroid 4-dehydrogenase activity" | 0.008 | 0.927 | 0.019 |
| **GO:0017176** | “phosphatidylinositol N-acetylglucosaminyltransferase activity" | 0.021 | 0.871 | 0.02 |
| **GO:0047429** | “nucleoside-triphosphate diphosphatase activity" | 0.138 | 0.926 | 0.023 |
| **GO:0042922** | “neuromedin U receptor binding" | 0.001 | 0.946 | 0.032 |
| **GO:0003810** | “protein-glutamine gamma-glutamyltransferase activity" | 0.011 | 0.802 | 0.136 |
| **GO:0042577** | “lipid phosphatase activity" | 0.027 | 0.873 | 0.179 |
| **GO:0004499** | “N N-dimethylaniline monooxygenase activity" | 0.075 | 0.893 | 0.213 |
| **GO:0008318** | “protein prenyltransferase activity" | 0.029 | 0.756 | 0.234 |
| **GO:0016209** | “antioxidant activity" | 0.716 | 0.832 | 0.258 |
| **GO:0004176** | “ATP-dependent peptidase activity" | 0.199 | 0.752 | 0.281 |
| **GO:0050660** | “flavin adenine dinucleotide binding" | 1.629 | 0.966 | 0.282 |
| **GO:0019955** | “cytokine binding" | 0.027 | 0.939 | 0.294 |
| **GO:0004713** | “protein tyrosine kinase activity" | 0.165 | 0.644 | 0.32 |
| **GO:0019838** | “growth factor binding" | 0.038 | 0.938 | 0.37 |
| **GO:0004044** | “amidophosphoribosyltransferase activity" | 0.031 | 0.869 | 0.438 |
| **GO:0004843** | “thiol-dependent deubiquitinase" | 0.209 | 0.752 | 0.491 |
